# Dynamic Transcription Factor Coupling Shapes Adaptive Enhancer Programs Driving Tumour Cell Heterogeneity and Growth

**DOI:** 10.64898/2026.11.24.769040

**Authors:** Sweta Sharma, Alireza Lorzadeh, George Ye, Moise Bonheur, Eric Sanchez, Juliana Valenzuela, Unmesh Jadhav

**Author notes:** **Corresponding author:** Unmesh Jadhav, PhD, Department of Stem Cell Biology and Regenerative Medicine, Keck School of Medicine, University of Southern California, Broad CIRM Center, 1425 San Pablo Street BCC-511, Los Angeles, CA 90033.

## Abstract

Tumour cells are inherently plastic and develop clonal heterogeneity early, laying the foundation for future invasion, metastasis, and resistance to therapy. The molecular basis of these early adaptations remains incompletely defined. Using inducible Apc deletion in mouse intestinal stem cells to mimic the most common initiating event in sporadic colorectal cancer (CRC), we show that cellular heterogeneity and adaptability in early adenomas reflect tumour-specific transcription factor (TF) activities that establish dynamic epigenetic programs. Enhancer profiles and single-cell epigenomes and transcriptomes reveal cooperative partnering between SOX17, which is selectively induced in adenomas, and constitutively active β-catenin. This cooperation activates adenoma-specific enhancers and generates clones with distinct gene-expression states that maintain stem cells in some populations and activate cell motility genes at the invasive front. These early epigenetic and cell states persist in distinct cellular clusters in advanced human CRCs. Loss of SOX17 attenuates target enhancers and associated genes, particularly those related to cell motility, and restricts adenoma growth. Accounting for incomplete adenoma eradication, however, other SOX17 target genes remain active by engaging alternative enhancers. Thus, epigenetic adaptability arises early in transformation, circumvents tumour cell vulnerability to TF perturbations, and supports cellular heterogeneity, plasticity, and malignant processes well into cancer progression. These findings have important implications for tumour evolution, treatment resistance, and approaches to develop new therapies.

---

Resistance to therapy and relapsed disease remain major problems in CRC, a leading cause of cancer deaths. Metastatic and therapy-resistant clones arise early in the disease course^1^, reflecting cellular heterogeneity and cell-intrinsic plasticity^2^. Tumour cells adopt a range of functional states, including stem-like populations that sustain growth and invasive cells that remodel tissue structure^3^. Heterogeneity also generates novel features and competing subclones, ensuring tumour survival even if some subclones are lost^4^. Although heterogeneity can arise from mutational variations, early human and mouse adenoma transcriptomes at single-cell resolution reveal substantial heterogeneity prior to mutational diversification^5–7^. While the microenvironment likely plays a role in tumour heterogeneity and adaptability, tumour cell-intrinsic mechanisms remain incompletely understood. At the level of individual cells, plasticity equips tumours to shift states in response to altered signaling or therapy, and epigenetic reprogramming is considered a key driver^8^, possibly through fetal gene reactivation^9,10^. However, the mechanisms by which tumour cells adopt and rewire fetal gene regulatory programs remain largely unknown. While human single-cell atlases capture a static endpoint of tumour heterogeneity^11^, they do not address how a single genetic initiating event, like *Apc* loss, dynamically alters a homeostatic stem cell and its enhancer landscape.

Here, we demonstrate that the diversity and adaptability of TF–enhancer regulatory modules serve as a shared foundation for both tumour cell heterogeneity and plasticity. Through integrated single-cell multiomic profiling, enhancer activity mapping, and TF-binding analyses, we show that even in a uniform Apc-loss background—the most common initiating mutation in sporadic colorectal cancer^12,13^—distinct TFs establish epigenetically heterogeneous cellular states. Furthermore, while recent work has shown that SOX17 promotes adenoma growth in established models^9^, the upstream mechanisms by which it interacts with core oncogenic drivers like the Wnt/β-catenin cascade remain completely unresolved. We find that transformation is driven by dynamic TF coupling, which includes the fetal TF SOX17 selectively induced in adenomas redirecting the homeostatic Wnt effector β-catenin to non-homeostatic targets, expanding enhancer usage and activating adenoma-specific programs. In particular, the SOX17 directed program regulates cell adhesion and motility processes, as its loss disrupts these pathways and limits adenoma progression. However, the enhancer network remains adaptable: transformed cells engage alternative enhancer regulatory modules to sustain expression of many genes upon SOX17 loss. Together, these findings reveal both the vulnerability and resilience of tumour-specific epigenetic states, highlighting how a flexible enhancer architecture underlies tumour cell heterogeneity, adaptability, and progression.

## RESULTS

### Epigenetic diversity drives early aberrant cell states in intestinal *Apc*-mutant adenomas

While cellular heterogeneity evolves as tumours progress^14^, single-cell RNA-seq studies in mice show that heterogeneity emerges immediately following acquisition of founding mutations such as those in *Apc*^6,7^. To investigate epithelial cell-intrinsic epigenetic remodeling as a possible basis for this heterogeneity, we used *Apc^Fl/Fl^*;*Lgr5^EGFP-CreER-T^*^2^;*Rosa26^LsL-tdTomato^* (hereafter *Apc^−/−^*) mice to disrupt *Apc* selectively in Lgr5^+^ intestinal stem cells (ISCs) (Fig. 1a, Extended Data Fig. 1a). Two tamoxifen injections constitutively activated Wnt signaling and induced hundreds of intestinal adenomas within 14 days (Extended Data Fig. 1a). Transformed cells within adenomas were labeled with tdTomato and, as expected, showed Lgr5^+^ ISC expansion and increased proliferation, evidenced by increased 5-bromo-2’-deoxyuridine (BrdU) incorporation (Extended Data Fig. 1a,b). To assess transcriptional and chromatin heterogeneity, we used fluorescence-activated cell sorting (FACS) to purify all tdTomato^+^ cells, including Lgr5-EGFP^+^ ISCs, from *Apc^−/−^* and wild-type (*WT*) mice for single-cell Multiome (scRNA-seq + scATAC-seq) analysis (10x Genomics, Extended Data Fig. 1c). After quality control using Seurat and Signac^15,16^, we retained 4,511 *WT* and 3,908 *Apc^−/−^* cells and integrated these data, with *WT* cells as the reference (Fig. 1b). Canonical markers identified *WT* cell populations (clusters), including *Lgr5^+^* stem and *Alpi^+^*enterocytes (Fig. 1c, Extended Data Fig. 1d). As expected, *Apc* loss disrupted differentiation, reducing enterocyte numbers^11,17^ (Fig. 1c,d, Extended Data Fig. 1d,e).

**Fig 1.**
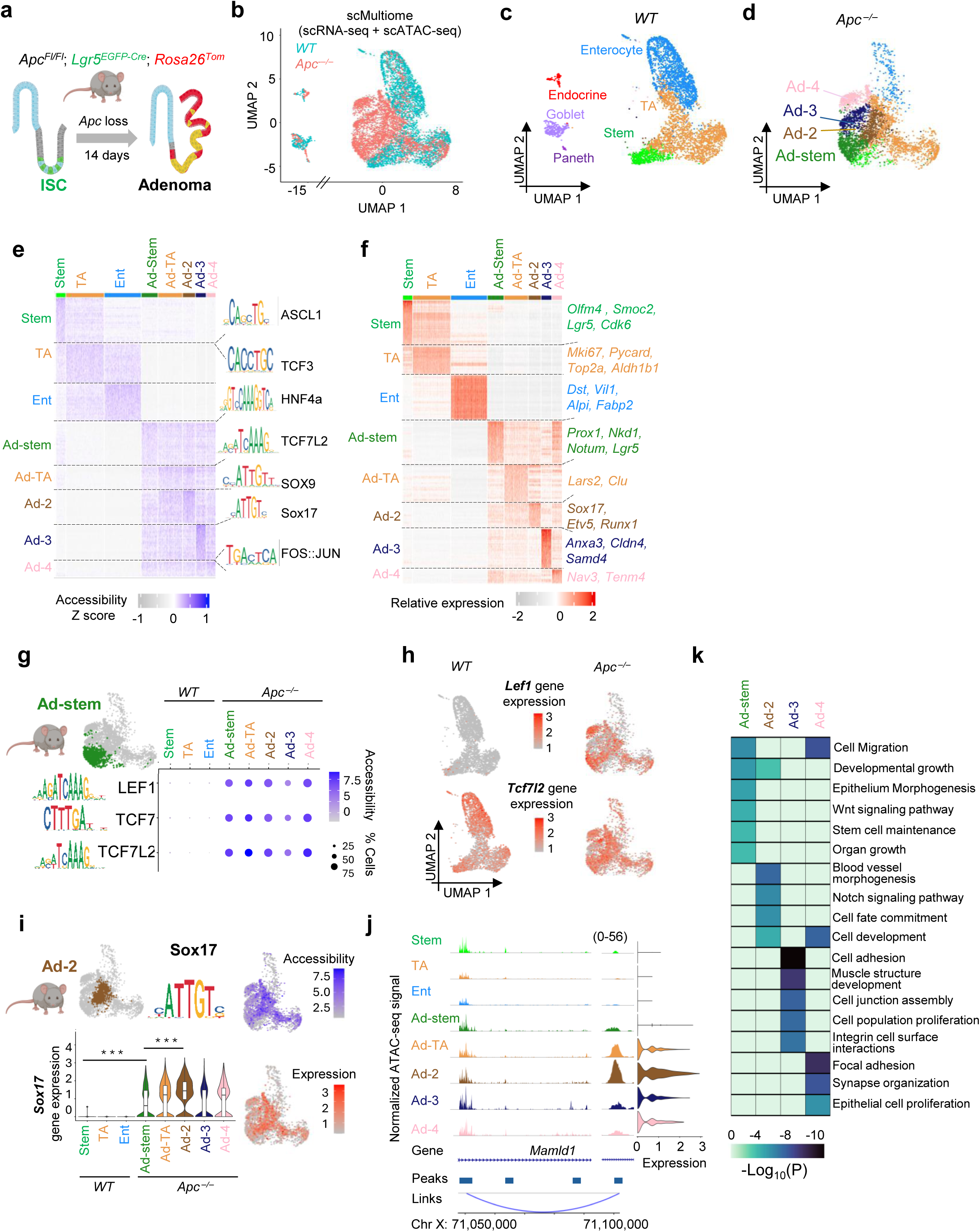
Single-cell multiome studies reveal diverse TF-enhancer modules driving cellular heterogeneity in intestinal adenomas. **a.** Schematic of the experimental mouse model demonstrating ISC transformation within 14 days following *Apc* deletion in adult mice aged ≥8 weeks and less than 6 months old. Wild-type Lgr5^+^ ISCs (green) are permanently labeled with tdTomato (red), enabling identification of transformed ISCs (yellow) and their progeny (red). **b.** UMAP representation of integrated scRNA-seq and scATAC-seq data for *WT* and *Apc^−/−^* cells. **c-d.** Split UMAP plot showing cell clusters identified in the *WT* sample (**c**) and *Apc^−/−^* cells (**d**). Colors indicate cell types. **e.** Heatmap showing relative accessibility of top 100 differentially accessible peaks in *WT* and *Apc^−/−^* cell clusters and significantly enriched MOTIFs (right) for each cluster (Fold enrichment > 1.86, *p* < 1.74e-89, hypergeometric test). **f.** Heatmap showing relative expression of top 100 differentially expressed genes per cluster (2-fold change and *padj < 0.05*, genes expressed in at least 10% of cells, Wilcoxon Rank Sum test). **g.** Dot plot showing gain of accessibility at *Ad-stem* specific loci with motifs of TCF/LEF factors in *Apc ^−/−^* cells compared to *WT* cells. Dot size represents the percentage of cells enriched for the TF motifs in *WT* and *Apc ^−/−^* cell clusters. **h.** Split UMAP of *WT* and *Apc^−/−^* cells showing expression of *Lef1* (Top) and *Tcf7l2* (Bottom). Heat indicates gene expression level. **i.** UMAP (red) and violin plot showing increased expression of Sox17 in adenoma cells. Cells in UMAP (blue) represent gain of accessibility at loci with enrichment of SOX17 motif in *Apc ^−/−^* cells. Wilcoxon test *p*-values are shown. (*p* < 0.001 = ***, *p* < 0.01 = **, *p* > 0.05 = NS). **j.** Genomic tracks showing normalized ATAC-seq signal (pseudobulk representation of scATAC-seq signal merged for all cells in each represented cluster) across *Mamld1* gene. Peaks (blue bars) are called using scATAC-seq data. Links in blue represent coordinated gain/loss of accessibility and gene expression across *WT* and *Apc ^−/−^* cells, supporting functional association of epigenetic change and gene expression. Violin plots on the side show expression of *Mamld1* gene in each cluster. **k.** Heatmap showing top enriched pathways for genes linked to *Ad-stem*, *Ad-2*, *Ad-3* and *Ad-4* specifically accessible peaks. Heat indicates FDR determined by Hypergeometric test corrected for corrected for multiple testing using the Benjamini-Hochberg algorithm.

Clustering of cells based on transcriptomic (scRNA-seq) and chromatin accessibility (scATAC-seq) identified *Apc^−/−^*-specific groups: adenoma-stem (*Ad-stem*), *Ad-2*, *Ad-3*, and *Ad-4*, each with distinct transcriptional and accessibility profiles (Fig. 1d, Extended Data Fig. 1d). The *Ad-stem* cluster in adenomas showed expansion of *Lgr5^+^* stem cells. Differential gene expression analysis revealed a robust, cell-intrinsic upregulation of *Nkd1 (8.69 fold change, padj = 4.36 x 10*^−130^), a known target of aberrant Wnt activation^18^ specifically within these adenoma associated stem cells compared to normal cells (Extended Data Fig. 1d). Notably, this induction was heterogeneous across individual adenoma stem cells rather than uniform across the expanded population, indicating that a common initiating mutation does not produce a uniform transcriptional response; while other clusters exhibited distinct gene signatures, including tumour growth associated genes such as *Cldn4*^19^ in *Ad-3*, indicating early emergence of multiple aberrant cell states (Extended Data Fig. 1d). Broad separation of *Apc^−/−^* and *WT* cells in ATAC Uniform Manifold Approximation and Projection (UMAP) space, compared to RNA-UMAP, was consistent with changes in chromatin accessibility preceding transcriptional diversification (Extended Data Fig. 1f,g). To examine this further, we quantified gains and losses of chromatin accessibility and gene expression in *WT* progenitor cells (TA and enterocytes) and *Ad-stem* relative to *WT* stem cells, identifying additional changes in *Ad-TA*, *Ad-2*, *Ad-3*, and *Ad-4* cells relative to *Ad-stem* (*q* <0.05, fold-change >1.2, Extended Data Fig. 1h,i). Differentially accessible peaks exceeded differentially expressed genes in number during *WT* stem cell differentiation (ISC → enterocytes) and changes in accessibility were most pronounced during ISC transformation upon Apc loss (Extended Data Fig. 1h,i). We note that this ordering is inferred from pseudotemporal relationships among cells collected at a single 14-day endpoint rather than measured over time. Because single-cell accessibility data are sparser and more binary than transcriptomic data, which can inflate apparent separation between cells, accessibility in these comparisons is quantified over pseudobulk-derived peaks shared across cell populations rather than over per-cell peak calls.

Pseudotime-based trajectories using Monocle ordered *WT* cells along clear trajectories, from stem cells to TA and absorptive (enterocyte) or secretory cells, and chromatin accessibility changes preceded transcriptional shifts (Extended Data Fig. 2a,b). Ordering wild-type cells along the transcriptional trajectory and scoring the same enterocyte genes in both modalities showed that 38.3% of cells in the earliest pseudotime bin already carried enterocyte-level chromatin accessibility at these genes, while 5.4% had reached enterocyte-level expression (Extended Data Fig. 2e-g). In contrast, none of the four *Apc^−/−^* clusters (*Ad-TA*, *Ad-2*, *Ad-3*, *Ad-4*) showed pseudotime transition from stem cells or aligned along a pseudotime axis (Extended Data Fig. 2c,d), suggesting rapid diversification of chromatin accessibility and transcriptional programs after *Apc* loss with distinct cell states that do not follow a conventional differentiation trajectory. In regions differentially accessible in *WT* cell clusters, ASCL1, TCF3, and HNF4A, known drivers of the intestinal lineage, were enriched (*padj* < 0.05) in *WT* stem, TA, and differentiated cells, respectively (Fig. 1e; Table 1). In contrast, *Ad-stem* cells were enriched for TCF and FOX motifs; *Ad-2* cells for SOX factors (including SOX17); and *Ad-3/4* clusters for AP-1 (FOS, JUN) motifs (Fig. 1e). These chromatin signatures mirrored unique transcriptional programs (Fig. 1f; Table 2), including transformation-associated (*Prox1*, *Nkd1*, *Clu*) and TF (*Sox17*, *Etv5*, *Runx1*) genes. Thus, even within a uniform *Apc*-mutant background, early adenomas diversify into distinct epigenetic and gene-expression states.

### Diverse TFs orchestrate early epigenetic rewiring in adenomas

We linked cluster-specific gains or losses of enhancer accessibility to expression changes of nearby genes using linkage analysis in Signac. This approach requires an accessibility change to coincide with a change in the expression of a nearby gene in the same cells, so it identifies regulatory changes that are already transcriptionally productive and excludes regions that gain or lose accessibility without a corresponding expression change. In the *Apc^−/−^*stem (*Ad-stem*) cells, gene-linked enhancers showed a marked gain in collective accessibility compared to *WT* stem cells, and this increase persisted across adenoma clusters (Extended Data Fig. 3a), indicating that substantial epigenetic remodeling during stem cell transformation (Extended Data Fig. 1i) persists in progeny cells. Because TFs influence chromatin remodeling and enhancer activity, we identified TFs whose motifs were enriched within the linked enhancers and whose expression was enriched in the same cell clusters, delineating functional, cluster-specific TF–enhancer–gene modules. The motif for β-catenin partners TCF and LEF^20–22^ was the most enriched at these enhancers in *Ad-stem* cells (Extended Data Fig. 3b, Table 1) and ChromVAR analysis confirmed increased accessibility at loci with enriched LEF1, TCF7, and TCF7l2 motifs in *Apc^−/−^*cells (Fig. 1g). *Lef1* was newly expressed in *Apc^−/−^* adenoma cells, suggesting TF expression–based control of its target enhancers (Fig. 1h, Extended Data Fig. 3c). In contrast, *Tcf7l2* was expressed in both *WT* and *Apc^−/−^* cells, but its target enhancers were activated only in *Apc^−/−^* cells, indicating adenoma-specific binding, likely mediated by a partner (Fig. 1h, Extended Data Fig. 3c). Thus, transformation-associated TF activation may enable different TF partnering events, redirecting β-catenin to new genomic targets and altering Wnt signaling outcomes and cell states. Genes linked to *Ad-stem*–specific TF–enhancer–gene modules were highly expressed in *Apc^−/−^* stem cells (Extended Data Fig. 3d, Table 3).

Enhancers uniquely accessible in *Ad-2* cells were enriched for SOX family motifs, particularly SOX9 and SOX17 (Extended Data Fig. 3e,f Table 1), TFs implicated in cell plasticity, developmental reprogramming, and immune evasion during early CRC initiation^7,9^. Notably, enhancers enriched for the SOX17 motif showed high accessibility alongside *Sox17* expression specifically in *Ad-2* cells (Fig. 1i). Genes linked to *Ad-2*–specific enhancers, including *Mamld1,* a known driver of tumourigenesis along with YAP1^23^, had elevated expression within this cluster (Fig. 1j, Extended Data Fig. 3g; Table 3). *Ad-3* and *Ad-4* clusters contained AP-1 (FOS/JUN)-motif-enriched loci (Extended Data Fig. 4a-f; Table 1), and invasion associated genes *Anxa3* and *Tenm4*^24^ linked to these loci showed elevated expression in the respective clusters (Extended Data Fig. 4g-i; Table 3).

Together, these findings indicate that distinct enhancer activation programs associated with TCF/LEF, SOX, and AP-1 families underlie heterogeneous epigenetic and transcriptional states in adenomas, with TCF/LEF–linked programs enriched for Wnt signaling, stem cell maintenance, and developmental processes, while SOX- and AP-1–associated programs for cell-fate determination, tissue morphogenesis, cell–cell adhesion, and epithelial organization (Fig. 1k).

### Molecular and cellular heterogeneity in early adenomas is preserved in advanced CRC

To ask whether transcriptional programs activated early in adenomas persist in fully developed human CRC, we analyzed RNA-seq data from The Cancer Genome Atlas (TCGA) Colon Adenocarcinoma (COAD) cohort of 454 tumours and 39 normal samples. Using the CMScaller algorithm for gene expression-based molecular classification (CMS1–4)^25^, we identified tumours belonging to the canonical Wnt-driven subtype CMS2 (Extended Data Fig. 5a). Within this group, 75–77% of genes upregulated in *Ad-stem*, *Ad-2*, *Ad-3*, and *Ad-4* clusters were significantly increased relative to normal tissue (DESeq2, *q* <0.05, Extended Data Fig. 5b). For example, FMNL2 and *LAMC2*, which promote tumourigenesis, angiogenesis^26^, and CRC invasion^27^, were upregulated in TCGA tumours and *Fmnl2* and *Lamc2* showed acquisition of new enhancers and elevated expression in *Apc^−/−^* mouse epithelial cells (Extended Data Fig. 5c,d). Thus, enhancer rewiring associated with TF activity established during early transformation persists into later stages of CRC, even when transient factors such as SOX17 are no longer expressed^9,28^.

To assess whether epigenetic and cell state heterogeneity established early in transformation is conserved during tumour progression, we analyzed published single-cell RNA-seq data from human CRC (23 tumour and 10 normal samples)^29^. In five independent CMS2 tumour samples comprising more than 1,000 epithelial cells, UMAP projections identified distinct *Apc^−/−^* mouse adenoma–defined clusters expressing *Ad-stem*, *Ad-2*, *Ad-3*, and *Ad-4* linked gene sets, highlighting persistence of early transformed cell states (Extended Data Fig. 5e). Moreover, mapping adenoma cluster–specific gene signatures (Extended Data Fig. 1d) to epithelial cells in human CRCs revealed retention of many signatures, particularly Wnt signal–linked genes in CMS2 tumours (Extended Data Fig. 5f). Accumulating driver mutations beyond *Apc* are expected to drive further chromatin remodeling during progression, but this has not been tested against enhancer programs defined at single-cell resolution. We therefore mapped bulk ATAC-seq from an isogenic organoid mutation series — *WT*, *Apc^−/−^* (A), *Apc^−/−^*;*Kras*^G12D^ (AK), *Apc^−/−^*;*Kras*^G12D^;*Trp53*^−/−^ (AKP) and *Apc^−/−^*;*Kras*^G12D^;*Trp53*^−/−^;*Smad4*^−/−^ (AKPS) intestinal stem cells^30^ — onto the Ad-stem, Ad-2, Ad-3 and Ad-4 enhancer sets (Extended Data Fig. 5g). Accessibility at all four sets increased significantly from WT to *Apc^−/−^* (*p* < 2.2 × 10^−16^ for each, two-sided Wilcoxon rank-sum test across enhancers), and this transition accounted for the largest shift in median signal at every set. Once opened, the Ad-3 and Ad-4 sets remained accessible across AK, AKP and AKPS, with Ad-4 maintaining a stable median signal throughout, and the multi-mutant genotypes showed no accessibility gain at regions outside those already established in *Apc^−/−^*cells. The enhancer spectrum that distinguishes adenoma subpopulations, including the Ad-3 and Ad-4 programs, is therefore configured upon *Apc* loss and is maintained rather than extended as other driver mutations accumulate. Together, these findings reveal that early TF-associated enhancer remodeling and cellular heterogeneity in *Apc*-mutant adenomas are established at initiation and are pervasive features of tumour evolution.

### Apc loss–induced transformation redirects β-catenin to novel genomic targets via developmental TF partners

APC mutations increase nuclear localization of the WNT signaling effector β-catenin and motifs for WNT-pathway TFs TCF/LEF are enriched at loci specifically accessible in human colorectal polyps and tumours^11,31^. In *Apc^−/−^* mouse tissues, we observed robust nuclear β-catenin 14 days post adenoma induction (Fig. 2a), in line with prior reports^31,32^, and single-cell multiome data revealed elevated TCF/LEF expression and increased accessibility at their target loci in adenoma stem cells (Fig. 1g,h). Because TCF/LEF family members recognize near-identical motifs, this enrichment reports engagement of the family as a whole rather than the identity of the factor bound at a given locus; differential use of individual TCF/LEF proteins to direct β-catenin to alternative genomic regions has recently been demonstrated^33^.

**Fig 2:**
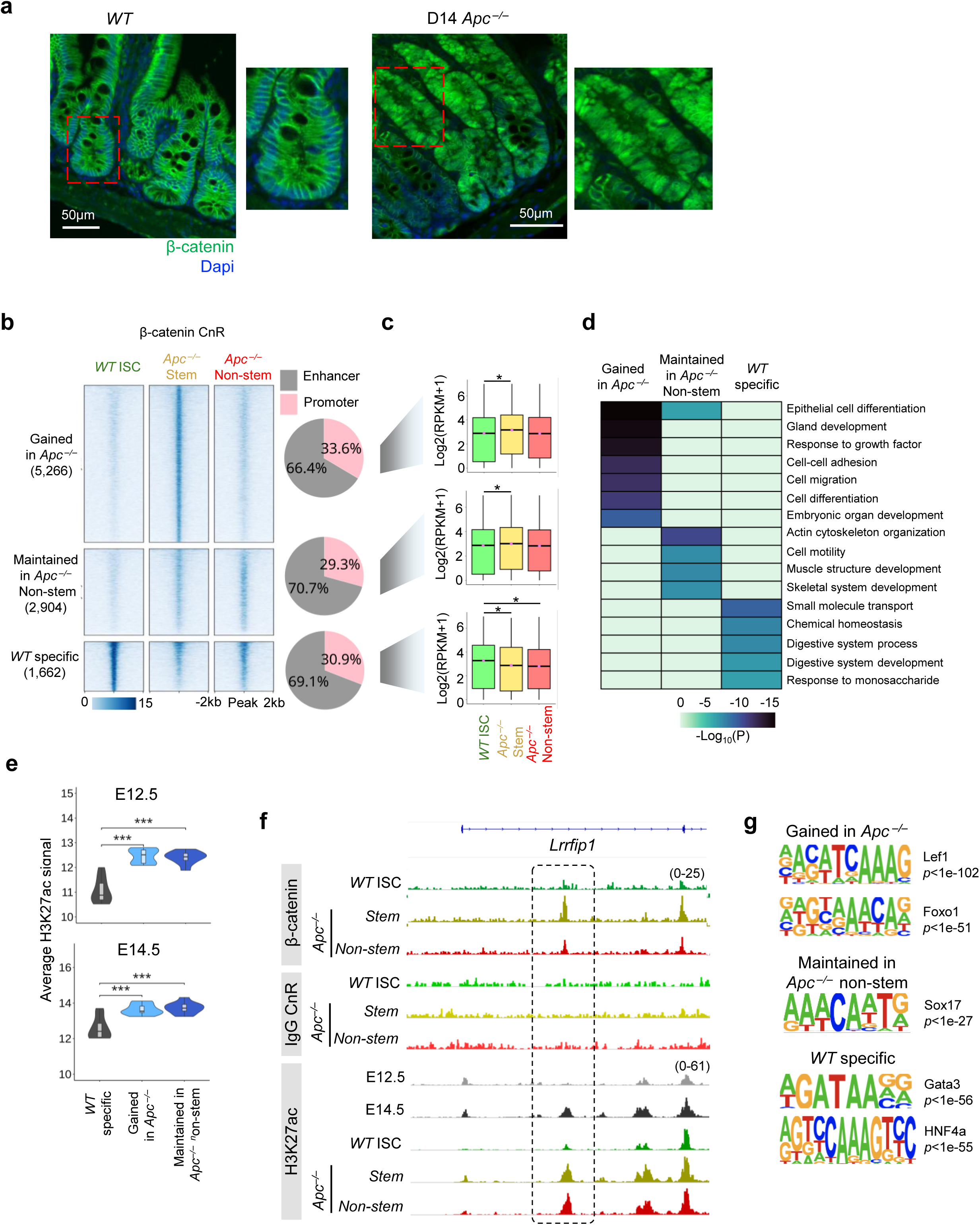
Non-canonical β-catenin targeting in adenomas. **a.** Fluorescent images showing β-catenin immunohistochemistry in mouse duodenum with DAPI labeling nuclei. Regions in red boxes are enlarged to show distribution of β-catenin in *WT* and adenoma cells from Day 14 *Apc ^−/−^* mice. **b.** Heatmap showing β-catenin CUT&RUN-seq (CnR) signal in a 2 kb window from peak centers at *WT* specific and *Apc ^−/−^* specific binding sites in *WT* ISC, and day 14 *Apc ^−/−^* stem and non-stem cells. Loci in each group are identified by k-means clustering of all peaks across the three cell types and represent significant signal change (*q* < 0.01). The matched IgG CUT&RUN control for these peak classes is shown in Extended Data Fig. 6a. Pie charts on the right show percentages of β-catenin binding sites in promoters (TSS −2 kb to +1 kb) and enhancers. **c.** Boxplots show gene expression (log2(RPKM+1)) of genes within 50 kb of the corresponding β-catenin bound enhancers (*p* < 0.05 two-sided t-test). Colors represent the indicated cell types. Black line within each box represents the median. **d.** Heatmap showing top enriched pathways for genes within 50 kb of β-catenin bound enhancers using Metascape analysis, ordered by significance shown in the heat scale. **e.** Violin plot showing quantification of H3K27ac signal in embryonic E12.5 and E14.5 small intestine^35^ for β-catenin-bound loci shown in **b.** Signal was quantified over 250 bp from peak centers, revealing increased developmental activity for novel β-catenin binding gained in *Apc ^−/−^*. *p* values were calculated using two-sided t-test. (*p* < 0.001 = ***, *p* < 0.01 = **, *p* > 0.05 = NS). **f.** Genome browser tracks showing β-catenin bound peaks (top) and enhancer mark H3K27ac (bottom), colored by cell type. Dotted box highlights enhancer active during development, deactivated in adult cells, and reactivated in *Apc ^−/−^* stem and non-stem cells. Matched IgG CUT&RUN tracks are shown for each genotype and cell population. **g.** Top Motifs enriched in three groups of β-catenin bound enhancers (in **b**) identified using HOMER analysis (*p* value are calculated using default binomial distribution).

Despite the known importance of altered Wnt/β-catenin signaling in transformation and tumour progression, the molecular details of β-catenin TF partner switching and its impact on genomic targeting and adenoma-specific enhancer activation remain unresolved. To address this, we first asked how β-catenin binding is altered in *Apc^−/−^* cells and conducted Cleavage Under Targets & Release Using Nuclease sequencing (CUT&RUN-seq) using FACS-sorted *Apc^−/−^* stem (Lgr5-EGFP^+^, tdTomato^+^) and non-stem (Lgr5-EGFP^−^, tdTomato^+^) cells (Extended Data Fig. 1a,c). K-means clustering using called peaks (MACS2^34^) showed widely remodeled β-catenin binding in *Apc^−/−^* cells (Fig. 2b); the matched IgG control showed no enrichment at these loci (Extended Data Fig. 6a). Notably, 5,266 loci gained β-catenin binding (1,767 promoters, 3,499 enhancers) specifically in *Apc^−/−^*stem cells, with reduced signal in non-stem cells. At 2,904 sites (851 promoters, 2,053 enhancers), β-catenin binding gained in *Apc^−/−^* stem cells was sustained in non-stem progeny. Conversely, β-catenin occupancy was reduced or lost at 1,662 loci (514 promoters, 1,149 enhancers). Expression of genes linked to β-catenin (within 50 kb) uniquely in *WT* ISCs was decreased in *Apc^−/−^* cells (Fig. 2c, Extended Data Fig. 6b). Conversely, genes linked to loci newly bound by β-catenin in *Apc^−/−^*stem (1,447) and non-stem (880) cells were significantly increased compared to *WT* cells, indicating that β-catenin redirection to non-physiologic enhancers activates adenoma-specific TF–enhancer–gene modules.

Enhancers bound by β-catenin in *WT* ISCs were enriched for digestion-related pathways, whereas those bound only in *Apc^−/−^* cells were enriched for developmental and cell migration processes (Fig. 2d). These loci also showed higher levels of histone mark H3K27ac, indicating elevated enhancer activity, in embryonic intestinal epithelium (E12.5–E14.5^35^) compared to *WT* ISC-specific enhancers (Fig. 2e), as exemplified by the *Lrrfip1* locus (Fig. 2f). Thus, although β-catenin is expressed throughout intestinal development and adulthood, its genomic occupancy appears tightly regulated and likely depends on epigenetic context and partner TFs. Motif analysis revealed enrichment of the TFs GATA3 and HNF4 at *WT*-specific β-catenin peaks (Fig. 2g). While transformed stem cells showed β-catenin association with TCF/LEF, sites with increased β-catenin binding in *Apc^−/−^* stem and non-stem cells were enriched for the SOX17 motif, suggesting a shift in TF partner landscape that accompanies redirection of β-catenin to adenoma-specific genomic targets.

### Sox17 supports adenoma growth and redirects β-catenin signaling

Given the enrichment of SOX motifs in *Ad2*-specific open chromatin peaks (Fig. 1i; Extended Data Fig. 3e,f) and SOX17 motifs at adenoma-specific β-catenin binding sites (Fig. 2g), we investigated SOX17 function in early adenoma formation. SOX17 is essential for definitive endoderm development^36^, where it is regulated by WNT–TCF signaling^37^ and directs β-catenin to developmental targets^38,39^; SOX17 also promotes carcinoma cell proliferation *in vitro*^40^. Normally absent in adult intestine, it is re-expressed in human colon polyps^9,28^ and mouse adenomas^41^ (Extended Data Fig. 6c) but silenced during CRC progression^9,42^, consistent with a transient role in early tumourigenesis. Supporting adenoma-specific reactivation, TCGA (454 tumours, 39 normals) and single-cell CRC datasets (23 tumours, 10 normals^29^) showed negligible *SOX17* expression in normal colonic epithelium or advanced tumours (Extended Data Fig. 6d,e).

In *Apc^−/−^* adenoma cells, where *Sox17* expression peaked in the *Ad-2* cluster together with motif enrichment (Fig. 1i; Extended Data Fig. 3f), we observed significant gains in chromatin accessibility and H3K27ac at multiple enhancers within 50 kb of the *Sox17* locus (Extended Data Fig. 6f). These enhancers are evolutionarily conserved and display increased chromatin accessibility, H3K27ac, and β-catenin/TCF7L1/SOX17 co-occupancy during human ES cell differentiation to definitive endoderm^39^ (Extended Data Fig. 6g). Together, these observations indicate reactivation of developmental enhancers accompanying *Sox17* expression in adenomas. Immunostaining confirmed SOX17 induction as early as the 7^th^ day after *Apc* deletion, extending by day 14 from Lgr5^+^ stem cells to tdTomato^+^ non-stem adenoma cells (Fig. 3a), while *WT* cells lacked any SOX17 staining, as expected (Extended Data Fig. 7a).

**Fig. 3.**
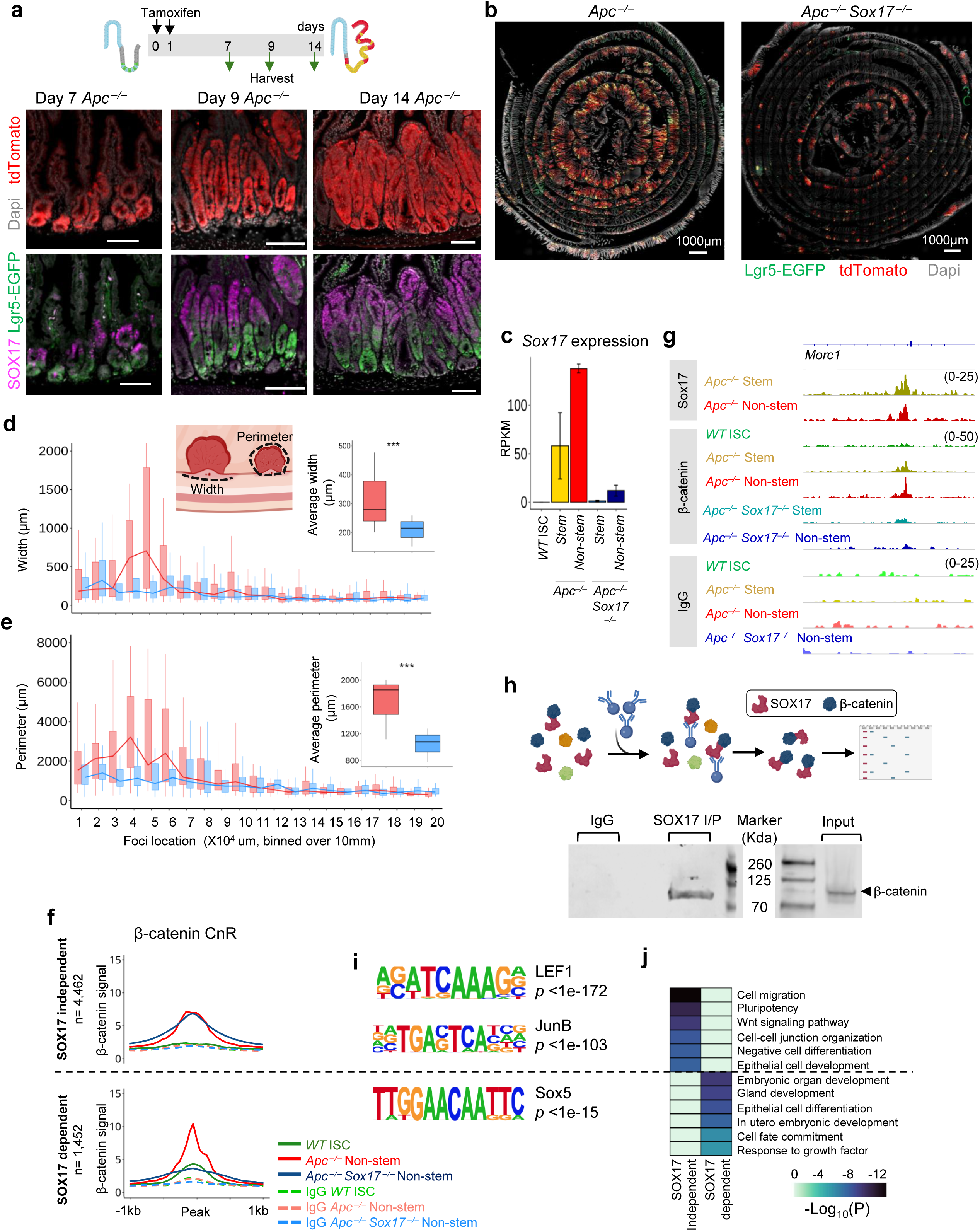
Loss of Sox17 impairs adenoma growth and alters β-catenin targeting. **a.** Fluorescent images showing immunohistochemical staining of tdTomato (all adenoma cells), SOX17, and Lgr5-EGFP (stem cells) in mouse duodenum sections harvested on Days 7, 9 and 14 post-*Apc* deletion, revealing increased Sox17 expression in adenomas. Co-staining with Lgr5-EGFP shows that Sox17 expression increases in Lgr5^−^ cells. Scale bar = 100 μm. **b.** Whole intestine harvested from day 14 *Apc ^−/−^* (left) and *Apc ^−/−^ Sox17 ^−/−^* (right) mice show overall tumour reduction. Scale bar = 1000 μm. **c.** Bar plot showing mRNA-seq expression of *Sox17* in WT ISCs, *Apc ^−/−^*, and *Apc ^−/−^ Sox17 ^−/−^* cells. Data are shown as mean ± SD, n = 2 replicates per condition. **d-e.** Illustration showing measurements used to obtain adenoma width and perimeter in *Apc ^−/−^*and *Apc ^−/−^ Sox17 ^−/−^*mice. Boxplots with trendline show the distribution of adenoma width and perimeter across the entire intestine (proximal to distal), binned every 10 mm of muscle length. Smaller box plots represent the average width and perimeter, demonstrating that adenomas in *Apc ^−/−^ Sox17 ^−/−^* are significantly smaller than in *Apc ^−/−^* mice. (*** *p* < 0.001, two-sided t-test) **f.** Profile plot showing β-catenin CUT&RUN-seq (CnR) signal at SOX17-dependent and independent enhancers (loci determined by k-means clustering of differential peaks with 2X fold change between any cell types, DEseq2, Wald test). Matched IgG CnR signal is plotted alongside for each genotype and cell population. **g.** Genome browser view showing a Sox17-dependent locus where β-catenin binding, present in *Apc ^−/−^* stem and non-stem cells, is reduced in *Apc ^−/−^ Sox17 ^−/−^* cells. Matched IgG CnR tracks are shown for each genotype and cell population. **h.** Western blot showing co-immunoprecipitation of β-catenin with SOX17 from *Apc ^−/−^* adenoma epithelial cell extracts. Adenoma cell lysates were immunoprecipitated with anti-SOX17 or with IgG and immunoblotted for β-catenin. Input, 10 percent of the material used per immunoprecipitation. β-catenin is recovered in the SOX17 immunoprecipitate and is absent from the IgG control. Representative of three independent experiments. **i.** Motif analysis on SOX17-dependent and -independent β-catenin bound regions using HOMER. Top enriched Motifs are shown for each peak set, with SOX-motif enrichment found in SOX17 dependent β-catenin targets. **j.** Heatmap showing top enriched pathways identified via Metascape for genes within 50 kb of Sox17-dependent and -independent β-catenin enhancers.

SOX17 CUT&RUN-seq in *Apc^−/−^* cells revealed 814 stem-specific binding sites, 1,500 sites gained in non-stem cells, and 1,012 shared enhancer sites, demonstrating engagement of distinct enhancer subsets across adenoma cell states (Extended Data Fig. 7b,c). SOX17-bound enhancers were highly accessible in *Apc^−/−^* cells but not in *WT* cells (Extended Data Fig. 7d), and were linked to genes involved in cell adhesion and motility, suggesting roles in epithelial reorganization and adenoma expansion (Extended Data Fig. 7d).

To examine SOX17’s role in redirecting Wnt/β-catenin activity and tissue remodeling, we generated *Apc^−/−^ Sox17^−/−^* double mutants by crossing *Apc^Fl/Fl^*;*Lgr5^EGFP-CreER-T^*^2^;*Rosa26^LsL-tdTomato^* and *Sox17^Fl^*(JAX 031712) mice (Fig. 3b, Extended Data Fig. 7e). mRNA-seq of FACS-purified Lgr5^+^ and Lgr5^−^ cells confirmed *Sox17* deletion (Fig. 3c), with rare escapees observed by IHC, a known feature of Cre-induced epithelial knockout^43^ (Extended Data Fig. 7f). Adenomas in *Apc^−/−^Sox17^−/−^* intestines were significantly smaller, with reduced basal spread and perimeter compared to *Apc^−/−^* controls (Fig. 3 b,d,e, Extended Data Fig. 7e), suggesting that SOX17 promotes adenoma growth.

β-catenin CUT&RUN-seq in stem and non-stem *Apc^−/−^ Sox17^−/−^* cells, compared with *WT* and *Apc^−/−^*samples, identified 4,462 loci where β-catenin binding persisted and 1,452 loci where binding was reduced in the absence of SOX17 (fold change ≥2, DESeq2) (Fig. 3f, g). Residual β-catenin signal at these loci remained above the matched IgG control, indicating reduced rather than abolished occupancy (Fig. 3f, Extended Data Fig. 7g); losses were most pronounced in Lgr5^−^non-stem cells, where *Sox17* expression is highest (Fig. 3a, c). In both enhancer groups, β-catenin recruitment was specific to adenomas and minimal in *WT* cells (Extended Data Fig. 7g). This β-catenin binding overlapped with SOX17 occupancy at the majority of these loci, suggesting that SOX17 facilitates β-catenin recruitment to noncanonical targets during transformation (Extended Data Fig. 7g). Consistent with this co-occupancy, SOX17 and β-catenin physically associate in adenoma cells: β-catenin was recovered in SOX17 immunoprecipitates from Apc−/−adenoma epithelial cell extracts and was absent from the IgG control (Fig. 3h). In further support of this point, sites that retained β-catenin binding after Sox17 loss (SOX17-independent) were enriched for TCF/LEF and JUN motifs, whereas those with reduced binding (SOX17-dependent) carried SOX motifs, closely similar to that for SOX17, alongside TCF/LEF motifs (Fig. 3i). β-catenin at these loci therefore engages SOX17 together with TCF/LEF rather than SOX17 alone. Both groups were linked to pathways involving Wnt signaling, MAPK cascades, cell junctions, and tissue development (Fig. 3j). Together, these findings indicate that early adenomas are characterized by a SOX17-associated shift in β-catenin genomic occupancy and implicate alternative TF partnering in enhancer choices and adenoma growth.

### Adenomas redistribute enhancer usage following *Sox17* loss

SOX17-dependent β-catenin targeting in adenomas, its heterogeneous expression in adenoma cells, and reduced yet persistent *Sox17^−/−^*adenomas together suggest that adenomas may adapt to SOX17 loss by remodeling enhancer usage. To identify such adaptation, we used H3K27ac native-ChIP-seq to profile enhancer activity in Lgr5-GFP^+^ (stem) and Lgr5-GFP^−^ (non-stem) cells from *Apc^−/−^ Sox17^−/−^*tissues alongside *WT* and *Apc^−/−^* controls. Differential H3K27ac analysis (fold change ≥2, *q* <0.05, DESeq2) and *k*-means clustering revealed extensive enhancer remodeling (Fig. 4a). Among altered sites, 7,757 enhancers lost activity upon *Sox17* deletion (*Sox17-*dependent), whereas 20,700 retained or gained activity (Sox17-independent): 11,161 sustained their activity, and 9,539 weakened in Apc*^−/−^* but strengthened in Apc*^−/−^ Sox17^−/−^* tissue. In contrast, 11,140 homeostatic enhancers were inactivated in adenomas, and as expected, these loci were enriched for motifs of known physiologic TFs such as HNF4A and KLF5^44^ (Extended Data Fig. 8a). *Sox17*-dependent enhancers showed highest accessibility and SOX motif enrichment in the *Ad-2* cluster, concordant with highest Sox17 expression in *Ad-*2 cluster (Fig. 4b, 1i), whereas *Sox17*-independent sustained and strengthened enhancers were enriched for FOS/JUN and FOSL2/ELF3 motifs, as observed in the *Ad-3* cluster (Fig. 4b, Extended Data Fig. 8b). These findings associate enhancer remodeling with distinct TF programs across adenoma subpopulations.

**Fig 4:**
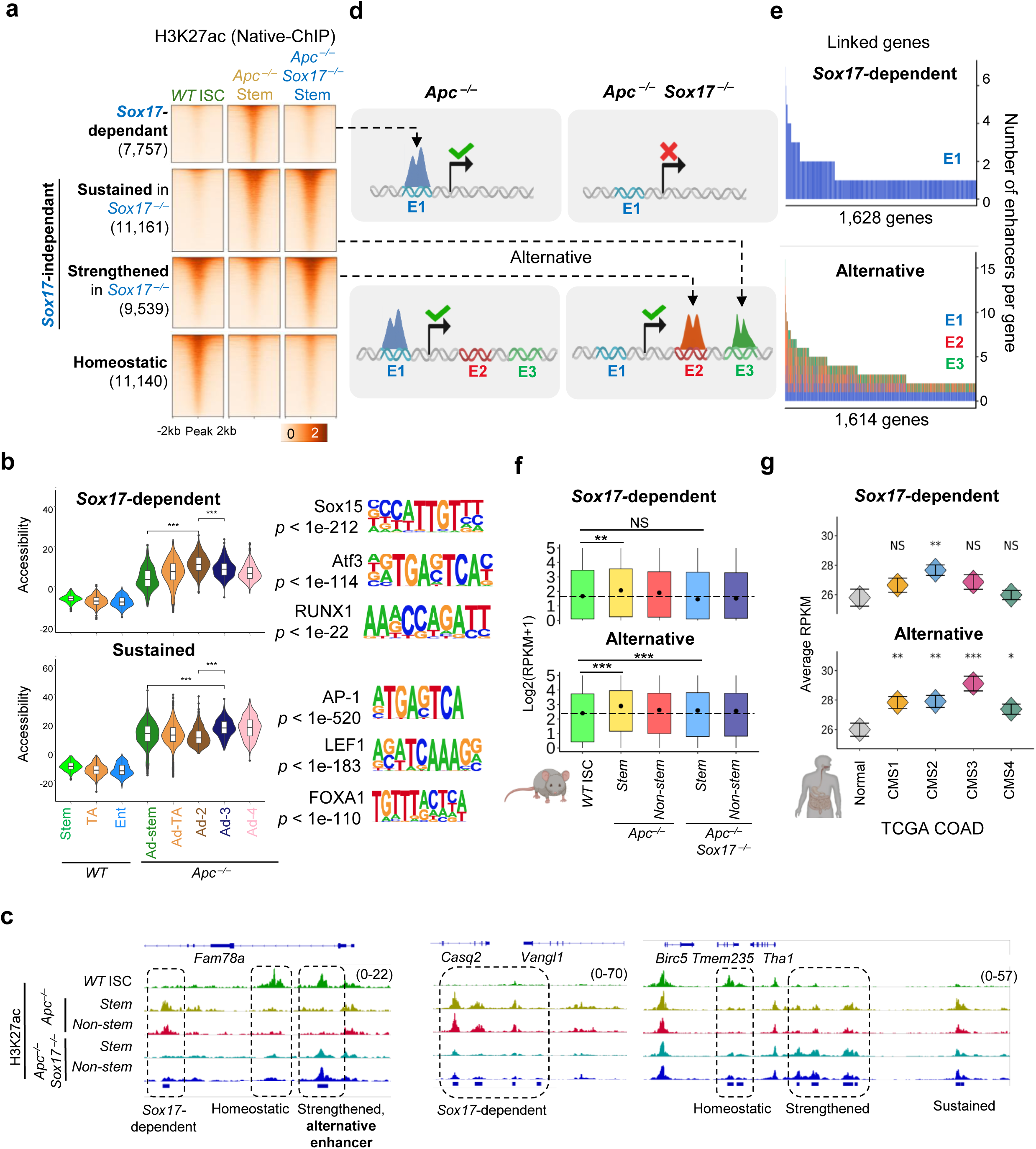
Enhancer adaptability upon loss of Sox17 in adenomas. **a.** Heatmap showing four clusters of H3K27ac enhancer peaks that are differential across the stem cells in *WT, Apc ^−/−^,* and *Apc ^−/−^ Sox17 ^−/−^* samples, with at least a 2-fold change in signal (*q < 0.05,* DEseq2 Wald test). **b.** Violin plot showing accessibility signal from scATAC-seq data in different *WT* and *Apc ^−/−^* clusters for enhancers lost in *Apc ^−/−^ Sox17 ^−/−^* (*Sox17*-dependent) and sustained enhancers from **a.** (*** *p* < 0.001, two-sided t-test). Top Motifs enriched for each group (HOMER) are displayed on the right. **c.** Genomic tracks highlighting genomic loci representative of the enhancer groups from panel **a**. **d.** Schematic showing engagement of alternative enhancers in absence of *Sox17* and deactivation of *Sox17*-dependent enhancers, which sustain the expression of the target gene. **e.** Bar plot quantifying the number of enhancers for genes that either have only *Sox17*-dependent enhancers or have alternative enhancers from other groups. **f.** Expression of genes linked only to Sox17-dependent enhancers (top, n = 1,628) and of genes that also have a redistributed enhancer (bottom, n = 1,614), in *WT* ISC and in stem and non-stem cells from *Apc^−/−^*and *Apc^−/−^ Sox17 ^−/−^* adenomas (Log2(RPKM+1)). Black dot indicates the median in each boxplot; the black dotted line marks the WT ISC median. Two-sided t-test: *** *p* < 0.001, ** *p* < 0.01, NS, not significant. **g.** Plot showing the average RPKM values for Sox17-dependent and alternative-enhancer genes *in TCGA-COAD.* (*p* < 0.001 = ***, *p* < 0.01 = **, *p* < 0.05 = *, *p* > 0.05 = NS, t-test)

To determine how SOX17 loss–associated enhancer changes affect gene expression, we examined transcriptional changes of genes linked to distinct enhancer groups in *Apc^−/−^* and *Apc^−/−^ Sox17^−/−^* stem (Lgr5^+^) and non-stem (Lgr5^−^) cells by mRNA-seq. In total, 7,062 genes were linked to these enhancers, of which nearly 20% were associated with multiple enhancers from different groups (Fig. 4c, Extended Data Fig. 8c). Notably, among the 3,242 genes linked to *Sox17*-dependent enhancers, ∼50% (1,614 genes) also harbored one or more nearby *Sox17*-independent (sustained or strengthened) sites, indicating redistributed enhancer engagement (Fig. 4c-e); the other 1,628 genes lacked such alternative enhancers. Genes with alternative enhancers maintained high expression in *Apc^−/−^ Sox17^−/−^*stem and non-stem cells, comparable to *Apc^−/−^* levels, whereas those without such enhancers downregulated to near–*WT* levels upon *Sox17* loss (Fig. 4f). In the TCGA-COAD dataset, genes linked to *Sox17*-dependent enhancers were selectively upregulated in CMS2 tumours, whereas genes associated with alternative enhancers were elevated across all CMS subtypes relative to *WT* colon, indicating persistence of adaptive enhancer programs in advanced CRC (Fig. 4g). Altogether, these findings indicate that *Sox17*-dependent programs preferentially engage genes linked to cell motility and tumour growth in CMS2 tumours, whereas loss of *Sox17* is accommodated through enhancer rewiring that sustains expression of genes broadly expressed across CRC molecular subtypes.

### Sox17 drives adenoma growth at the invasive front

To define the molecular basis of reduced adenoma size and spread in *Apc^−/−^ Sox17^−/−^* mice (Fig. 3d,e), we identified genes linked to *Sox17*-dependent enhancers (Fig. 4a) using scMultiome-based enhancer–gene linkage analysis, revealing 695 genes with concordant changes in enhancer accessibility and expression during transformation (within 50 kb of transcription start sites). These genes were enriched for pathways related to cell-cell adhesion, cell motility, and tissue morphogenesis (Fig. 5a), suggesting that SOX17 loss may impair epithelial structure and influence adenoma growth. Most genes in these pathways were strongly upregulated in *Apc^−/−^*, but reduced in *Apc^−/−^ Sox17^−/−^*, stem and non-stem cells (Extended Data Fig. 9a). This included cadherins and collagens such as *Cdh5* and *Col4a1*, which were activated in *Apc^−/−^* cells with SOX17 binding at nearby enhancers, but both the enhancer activity and expression were lost in the absence of SOX17 in *Apc^−/−^ Sox17^−/−^*cells (Fig. 5b,c, Extended Data Fig. 9b), revealing dependance of these enhancers and genes on SOX17’s action.

**Fig 5:**
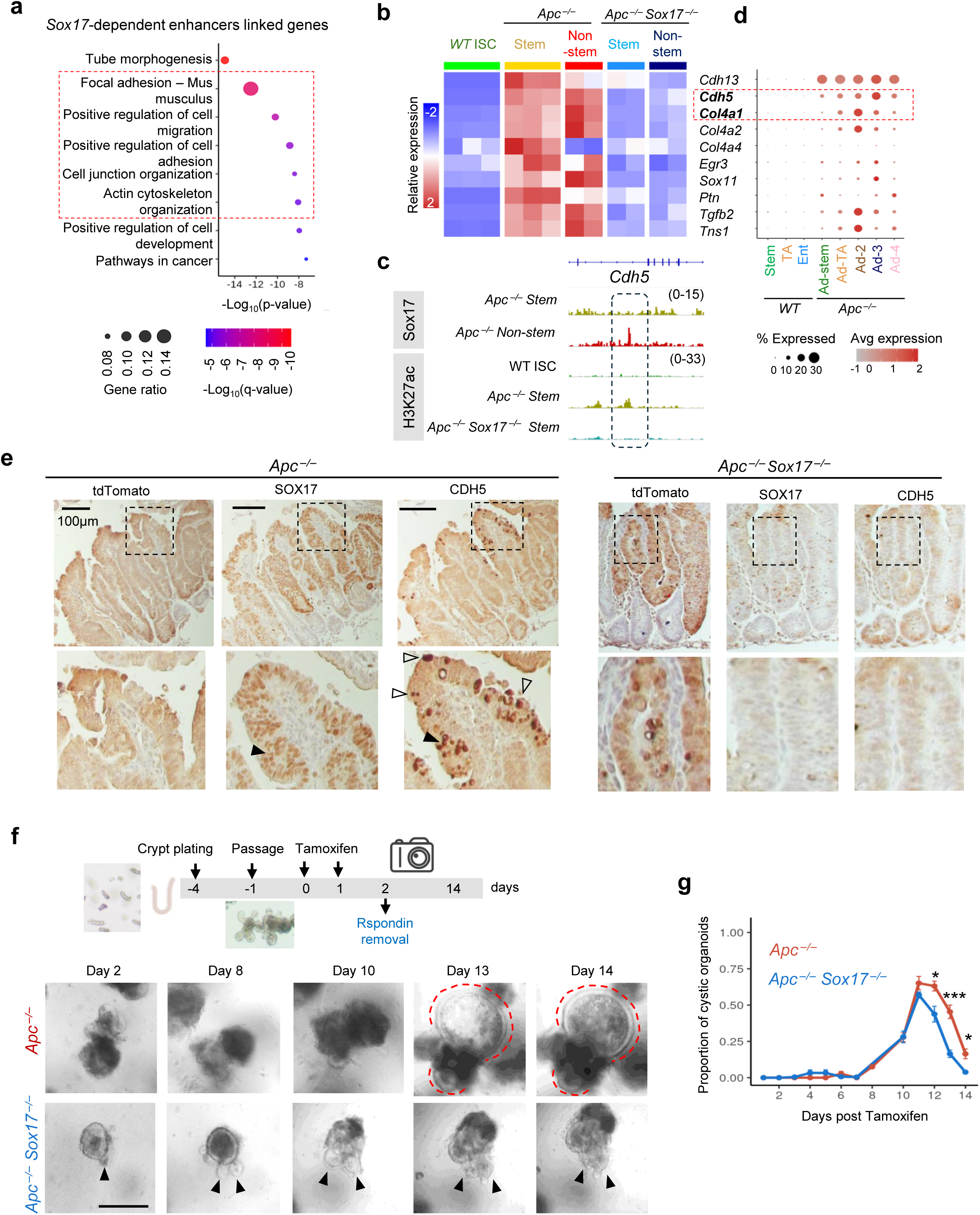
Sox17 controls structural growth of adenomas. **a.** Genes linked to *Sox17*-dependent enhancers from Fig. 4a in the scMultiome data from *WT* and *Apc ^−/−^* show enrichment of pathways regulating cell motility and cell adhesion (red dotted box). Heat indicates FDR determined by Hypergeometric test corrected for multiple testing using the Benjamini-Hochberg algorithm; dot size represents gene ratio. **b.** Expression of selected cell adhesion pathway genes from panel (**a**), shown as relative Log2(RPKM+1) across *WT, Apc ^−/−^,* and *Apc ^−/−^ Sox17 ^−/−^* stem and non-stem cells. **c.** Genome browser view of the *Cdh5* locus showing active enhancers (H3K27ac) signals and SOX17 CnR peaks. Dotted box highlights an enhancer bound by SOX17 in *Apc ^−/−^* non-stem cells that loses its activity upon loss of Sox17. Matched IgG CnR tracks are shown for each genotype and cell population. **d.** Dot plot showing scRNA-seq expression of selected focal-adhesion pathway genes from panel (**a**) in *WT* and *Apc^−/−^*cells, highlighting the upregulation of *Cdh5* and *Col4a1 (*red dotted box) in *Ad-2* and *Ad-3* clusters distinct from stem cells. Dot size represents percentage of cells expressing the gene. **e.** Brightfield images showing IHC staining for SOX17, CDH5, and tdTomato in serial section using DAB staining. Black dotted boxes show enlarged regions where SOX17 and CDH5 co-expression is observed in *Apc ^−/−^* mice, whereas CDH5 expression is lost upon SOX17 deletion in adenomas. Arrowheads with black fill indicate nuclear SOX17 staining with corresponding Cdh5 expression, whereas open arrowheads indicate peripheral and non-nuclear CDH5 staining. **f.** Schematic (top) represents the experimental timeline for the organoid assay. Crypts harvested from *Apc^−/−^,* and *Apc^−/−^ Sox17 ^−/−^* mice were cultured for 5 days before Tamoxifen addition to delete *Apc* and *Sox17* in culture. Brightfield images (below) show structural changes post-selection and formation of cystic organoids at selected time points. Red dotted lines represent cystic structure in organoids and black arrow heads show outgrowths that fail to grow cystic structure in *Apc ^−/−^Sox17 ^−/−^.* Scale bar = 50 μm. **g.** Line plot showing the proportion of cystic organoids among all organoids at the indicated days. Significance between *Apc ^−/−^,* and *Apc ^−/−^ Sox17 ^−/−^* is shown (*p* < 0.001 = ***, *p* < 0.01 = **, *p* < 0.05 = *, two-sided t-test, n = 3 mice per condition).

Single cell transcriptomic data revealed that although *Cdh5* and *Col4a1* were activated in *Apc^−/−^* stem cells, their expression peaked in *Ad-3* and *Ad-2* clusters, respectively (Fig. 5d), suggesting SOX17’s role beyond adenoma stem cells. As growing adenomas exhibit increased cell motility at the peripheral growth (invasion) front^45^, we tested the spatial specificity of CDH5 in *Apc^−/−^* and *Apc^−/−^ Sox17^−/−^* adenomas. Immunohistochemistry revealed spatially restricted CDH5 at adenoma peripheries distal to stem cell–rich cores, where SOX17 was also high (Fig. 5e, 3a). Indeed, CDH5 was present in the cytoplasm of cells with nuclear SOX17 and occasionally in SOX17⁺ nuclei. In *Apc^−/−^ Sox17^−/−^* adenomas, CDH5 was largely lost, with sporadic expression in rare escaper cells(Fig. 5e, Extended Data Fig. 7f). These data suggest a role of SOX17 activity at adenoma peripheries in remodeling epithelial structure and promoting adenoma growth and spread, whereas SOX17 loss restricts these processes.

To test epithelial-intrinsic SOX17 requirements, we generated *Apc^−/−^*and *Apc^−/−^ Sox17^−/−^*organoids from mouse crypts, induced gene deletion with tamoxifen, and monitored structural and growth changes over 14 days (Fig. 5f). As expected, *Apc^−/−^* organoids formed cystic structures indicative of transformation^46^, whereas significantly fewer *Apc^−/−^ Sox17^−/−^* organoids formed or maintained cystic structures (Fig. 5f,g, Extended Data Fig. 9c), supporting SOX17’s role in epithelial structural remodeling and adenoma expansion. In line with SOX17’s essential function in modulation of fetal endodermal structure^36,39^, other SOX17-targeted cell adhesion and motility genes identified in adenomas (Fig. 5b) were also active in SOX17+ fetal endodermal cells^47^ (Extended Data Fig. 10a-c). Collectively, these findings reveal that SOX17-dependent enhancer rewiring reactivates fetal tissue remodeling programs during early tumourigenesis, thereby constructing a flexible epithelial regulatory architecture that sustains heterogeneity and drives adenoma progression.

## DISCUSSION

TFs act as the terminal effectors of cell-extrinsic signaling, translating pathway activity into gene expression. Wnt signaling plays a central role across tissues by regulating stem cell identity, tissue architecture, and growth, and its outputs depend on precisely controlled TF partnerships to activate spatiotemporally restricted targets^48^. β-catenin, the core Wnt effector, is regulated through nuclear shuttling and interactions with specific TFs that direct its genomic binding and define Wnt outcomes. Certain TCF/LEF family TFs and SOX17 are specifically expressed during mouse and human fetal development and activate Wnt target genes required for cell motility and tissue morphogenesis^39^. How this TF-directed routing of Wnt signaling is repurposed during colorectal tumourigenesis, and the direct targets of β-catenin in transformation have remained unclear. Our findings show that partner-TF availability represents a key regulatory node that constrains transformation. In adulthood, silencing fetal-restricted TFs restricts β-catenin to adult-specific targets, limiting inappropriate Wnt outputs even under constitutive β-catenin activation. In contrast, tumours reactivate fetal TFs, enabling β-catenin to access developmental programs that support growth and transformation.

Tumour heterogeneity, characterized by coexisting clones with distinct transcriptional and chromatin states, is essential for cancer progression and relapse. Such diversity enables tumours to withstand environmental pressures, as loss or failure of one clone can be offset by persistence of others, sustaining overall growth. While cancer-type-specific epigenetic wiring is thought to underlie the heterogeneity^49^, its origins have remained incompletely understood. Our data indicate that heterogeneity arises from differential TF activation across tumour cells, enabling tumours to meet diverse ecological and structural demands. Specialized TF-driven regulatory networks thus provide a blueprint for molecular diversification among tumour cells. Human colorectal polyps are enriched for TCF/LEF TFs^11^, and we show that SOX17 reroutes β-catenin to TCF/LEF-motif-enriched loci in early adenomas, increasing chromatin accessibility and activating target gene expression. These findings illustrate how combinatorial Wnt–partner TF interactions function differently in fetal versus adult contexts and how tumours exploit this circuitry. Deletion of *Sox17* blocks β-catenin rerouting and tumour-specific gene activation, highlighting partner TFs and their associated epigenetic modules as actionable vulnerabilities. Prior work on Fos/AP-1–dependent fetal gene activation^10^ and Sox9-mediated fetal reprogramming^7^ further supports TF-driven rewiring as a common oncogenic strategy. Among the Wnt effectors, LEF1 is the only TCF/LEF family member absent from normal adult intestinal epithelium and induced during intestinal tumourigenesis^21^, mirroring the developmentally restricted expression of SOX17 itself^36–38^. AP-1 factors, by contrast, are constitutively expressed in intestinal epithelium and acquire context-specific binding in injury and disease. Adenoma-specific enhancer configurations may therefore arise through two routes: re-engagement of developmentally restricted factors, and redirection of constitutively expressed ones. We test the first directly, through SOX17. Importantly, our data reveal that this early transformation window represents a highly dynamic process rather than the endpoints captured by human tissue atlases^11^. The TCF/LEF- and AP-1-associated states we identify are defined by their chromatin and expression signatures rather than by loss of function, so TF coupling as a driver of heterogeneity is exemplified here by SOX17. By resolving how SOX17 redirects β-catenin to non-homeostatic genomic targets, our study supplies a molecular mechanism missing from prior tumour models. In addition, our data show that these TF-driven programs are deployed heterogeneously rather than uniformly across tumour cells.

Spatial specialization further shapes tumour organization. Wnt-regulated genes are enriched in the stem-cell-rich tumour core, whereas the invasive edge is immunosuppressive and may depend on AP-1 and NF-κB activity^45^. Recent literature highlights the involvement of SOX17 in proliferation, epithelial-immune microenvironment crosstalk and immune evasion, as *Sox17*-deficient adenomas exhibit increased immune cell infiltration^9^. Here we reveal that SOX17 drives expression of numerous cell-motility genes through their enhancers at the invasion front, and that loss of SOX17 disrupts this program and limits tumour progression. These findings indicate that distinct TF targeting across tumour compartments supports acquisition of specialized properties required for growth and invasion. Tumour heterogeneity therefore reflects not only cell-intrinsic molecular evolution or microenvironmental influence, but coordinated activation of TF–enhancer modules that meet spatially distinct tumour demands and enable structural reorganization beyond normal epithelial constraints.

Beyond heterogeneity, cell-intrinsic plasticity enables transitions between tumour states that drive metastasis and relapse. Metastatic CRC lesions often share the same mutations as primary tumours, underscoring the role of TF-based reprogramming and enhancer rewiring rather than new genetic alterations. In CRC, metastasis frequently involves Lgr5⁻ cells that regenerate Lgr5⁺ stem cells at secondary sites and restore tumours following Lgr5⁺ cell ablation^3,50–52^, highlighting the need to target plasticity mechanisms. Here we identify chromatin circuits that remain flexible even as TF dependencies shift. Enhancer networks sustaining key growth-linked genes can switch to alternative regulatory elements when primary enhancers fail, such as following *Sox17* loss. This association is supported by the contrast between genes that retain a nearby Sox17-independent enhancer, which sustain expression after Sox17 loss, and those that do not, which fall to near-WT levels. Whether these redistributed enhancers are themselves required for adenoma persistence remains to be tested. Because each gene engages its own alternative element, requirement must be established enhancer by enhancer, which across the 1,614 genes concerned defines a separate line of investigation. Thus, tumour adaptability emerges from the combined effects of cellular heterogeneity and enhancer-system plasticity. Moreover, early-acquired epigenetic programs persist throughout CRC progression^11^, and we reveal that the regulatory states established in early adenomas are maintained in advanced tumours. Because SOX17 is silenced as tumours progress, the durable object of interest is not the factor itself but the regulatory logic it establishes. The enhancer programs it sets up are formed at initiation, persist after its expression is lost, and act on cell adhesion and motility genes, most prominently at the invasive front. Adhesion and motility are required not only for adenoma formation but through tumour expansion and invasion, so an early epigenetic event continues to govern a property relevant at every subsequent stage. Our data define this logic end to end, from the partner TFs that establish these enhancers, through the enhancer rewiring that sustains them when a founding factor is lost, to the target genes they control, such as *Cdh5*. Targeting TF–enhancer mechanisms that support both heterogeneity and plasticity may therefore provide strategies to limit relapse and improve patient outcomes.

## MATERIALS and METHODS

### Experimental model and subject details

*ROSA26^LSL-tdTomato^* and *Sox17^Fl^* (JAX# 031712), and *Lgr^EGFP-IRES-CreERT^*^2^ (JAX# 008875) mice^53^ were purchased from the Jackson laboratories. *Apc^fl^* mice were a gift from C. Perret^54^. To induce intestinal deletion of *Apc*, *Apc^Fl/Fl^* animals were bred with *Lgr5*^EGFP-IRES-CreERT2^ mice. This line was then crossed with *ROSA26^LSL-tdTomato^*to allow conditional fluorescent tdTomato labeling of Lgr5^+^ stem cells and their descendant populations. For targeted removal of *Sox17*, the colony was additionally intercrossed with *Sox17^Fl^* mice. Verification of engineered alleles was performed using PCR-based genotyping both at weaning and at experimental timepoints. Animals were housed at 23 ± 1°C with 50 ± 10% humidity under a 12-hour light/dark cycle, and were maintained according to protocols approved by the University of Southern California Department of Animal Resources. Experimental subjects showing clinical concerns — including decreased mobility, body weight loss exceeding 20%, hunched posture, or ruffled fur — were excluded and euthanized. Terminal CO2 exposure lasted 5 minutes, using a delivery rate equivalent to ∼20% of chamber volume per minute, following the 2013 AVMA Guidelines for Euthanasia of Animals. Cervical dislocation was applied as a confirmatory method.

### Mouse treatments

To trigger *Apc* or *Sox17* deletion, adult mice aged ≥8 weeks received 2 mg tamoxifen via intraperitoneal (i.p.) injection on two back-to-back days, with tissues collected at the specific time points designated for each experiment. Maintenance tamoxifen injections (alternate day after initial doses) were used for *Apc* and *Sox17* double deletion. To assess cell proliferation, mice were injected i.p. with 1 mg BrdU in PBS, and intestinal tissue was collected 1 hour later.

### Isolation of intestinal epithelial cells

The proximal one-third of the intestine (duodenum) from adult (≥8 weeks and less than 6 months old) *Apc^Fl/Fl^*; *Lgr5^EGFP-CreER-T^*^2^; *Rosa26^LsL-tdTomato^ (Apc^−/−^)* and *Apc^Fl/Fl^*; *Sox17^Fl/Fl^*; *Lgr5^EGFP-CreER-T^*^2^; *Rosa26^LsL-tdTomato^ (Apc^−/−^ Sox17^−/−^)* male and female mice were used for all genomics experiments. Freshly harvested intestines were washed in ice-cold PBS and incubated in 5 mM EDTA in PBS (pH 8) at 4°C for 40 min with vigorous shaking every 10 min. The released crypt epithelium was filtered through 70 µm strainers (BD Falcon). Crypts in the filtrate were dissociated into single cells using 4% TrypLE (ThermoFisher #A1217702) in DMEM at 37°C for 30 min, quenched with an equal volume of DMEM, and pelleted by centrifugation. Viable (DAPI-) EGFP^+^ tdTomato^+^ and EGFP^−^ and tdTomato^+^ cells were isolated by flow cytometry on a BD SORP FACSymphony S6 instrument.

### Mouse intestinal organoid culture

Mouse intestinal organoid cultures were prepared according to previously reported methods^55^. All steps for organoid derivation occurred on ice or at 4°C unless a different temperature was noted. Organoids were generated from *Apc^−/−^* and *Apc^−/−^ Sox17^−/−^*mice. Briefly, approximately 10 cm segments of proximal duodenum were excised, rinsed using PBS, and cut into pieces about 5 mm long. These fragments underwent incubation in 2 mM EDTA on a shaker at 4°C for a duration of 40 min, then were resuspended in Advanced DMEM/F12 and shaken vigorously. The resulting mixture was filtered using a 70-μm strainer and washed three times with medium to eliminate remaining EDTA.

Isolated crypts were collected by centrifugation at 300g for 5 min at 4°C, and 1 μl of the pellet was combined with 50 μl Matrigel (Corning, #356231), yielding 50–100 crypts in each Matrigel droplet. These droplets were dispensed into 24-well plates, allowed to solidify at 37°C, and then covered with mouse ENR medium (small intestine formulation) supplemented with ROCK inhibitor (Y-27632, 10 μM, Stemcell Technologies #72304). Mouse ENR included 10% R-spondin-conditioned media derived from 293T cells expressing murine R-spondin1, murine EGF (50 ng/mL, Peprotech #315-09), murine Noggin (100 ng/mL, Peprotech #250-38), plus 10 mM HEPES (Thermofisher #15630080), 1X Glutamax (Thermofisher #35050061), 1X B27 (Thermofisher #17504044), 1X N2 (Thermofisher #17502048), 1.25 mM N-Acetyl-Cysteine (Sigma #A8199), and Primocin and Normocin (Invivogen #ant-pm-2 and #ant-nr-2). For routine propagation, cultures were passaged every 3–4 days by resuspension in Cell Recovery Solution (Corning #354253) followed by centrifugation (300g, 5 min, 4°C), and organoids were replated in fresh Matrigel. Medium was renewed every 2 days.

Induced deletion of Apc and Sox17 in vitro was achieved by introducing 0.5 μM 4-OH tamoxifen (Sigma #H6278) to the culture medium for two days. Two days after the start of tamoxifen exposure, both ROCK inhibitor and R-spondin-conditioned medium were excluded to enrich for organoids lacking *Apc* or lacking both *Apc* and *Sox17*. Cultures continued for 14 days, with media changes every 48 hr. Organoids that expressed endogenous EGFP and tdTomato were imaged live using a KEYENCE BZ-X800 microscope.

### Immunofluorescence and IHC

Mouse intestines were fixed in 4% paraformaldehyde (EMS #15714) overnight, rinsed with PBS, and placed in 70% ethanol for storage. For immunohistochemistry, 5 μm paraffin sections were deparaffinized and rehydrated, followed by antigen retrieval in 10 mM sodium citrate buffer (pH 6) using heat and pressure. Sections were blocked in 2% BSA (Genesee #25-529) containing 0.1% Triton X-100 (Sigma #T8787) to reduce nonspecific antibody binding. For peroxidase-based detection, endogenous peroxidase was quenched using 3% hydrogen peroxide in methanol.

Slides were then incubated overnight at 4°C with antibodies against Sox17 (R&D Systems #AF1924), GFP to detect Lgr5-EGFP (Santa Cruz Biotechnology #sc-9996), RFP for tdTomato-expressing cells (Rockland #600-401-379), BrdU (Abcam #ab6326), Cdh5 (Proteintech #27956-1-AP), or β-catenin (Thermofisher #71-2700), each diluted in 1% BSA containing 0.1% Triton X-100. Biotin-conjugated anti-mouse, anti-goat, anti-rabbit, or anti-rat IgG (Jackson Laboratories, 1:1000) were used for detection, followed by Vectastain Elite ABC Kit (Vector Laboratories #PK-6100) and DAB Substrate Kit (Abcam #ab64238). Serial tissue sections were examined to determine Sox17+, tdTomato+, and Cdh5+ cell localization. For colocalization experiments, antigen-retrieved sections were stained with combinations of Sox17, BrdU, RFP, and/or GFP antibodies, followed by Alexa Fluor 488, 546, or 647-conjugated secondary antibodies raised in goat, rabbit, rat, or mouse. Slides were mounted using antifade medium with DAPI (Vector Laboratories #H-1200-10) to visualize nuclei.

For lineage tracing of Lgr5-EGFP intestinal stem cells in *Apc^−/−^*and *Apc^−/−^ Sox17^−/−^* mice, intestines were fixed, embedded in OCT compound (Tissue-Tek #4583), and flash-frozen. Fluorescent microscopy was performed on 7-μm sections to assess marker expression patterns.

### Single-cell multiomics sequencing

Single-cell multiomic libraries were generated following the recommended workflow for the Chromium Next GEM Single Cell Multiome ATAC plus Gene Expression Kit (10x Genomics #PN-1000285). In summary, nuclei were extracted from FACS-purified cells obtained from *WT and Apc^−/−^* samples. The resulting nuclei suspensions from both genotypes were loaded onto a Chromium Next GEM Chip J to form GEMs and apply unique molecular barcodes. Approximately 10,000 nuclei were targeted per sample, and parallel libraries for scRNA-seq as well as scATAC-seq were produced using the manufacturer’s chemistry. Sequencing of scRNA-seq libraries was performed to a depth of 20,000 reads for each nucleus, while scATAC-seq libraries were processed targeting 25,000 reads per nucleus, using the Illumina NovaSeq 6000 sequencing system.

### RNA isolation and bulk RNA sequencing

For bulk cell analyses, EGFP^−^ tdTomato^+^ or EGFP^+^ tdTomato^+^ populations were sorted using a BD SORP FACSymphony S6 flow cytometer, and the resulting cell pellets were flash frozen. RNA was extracted with the mirVana RNA isolation kit (Applied Biosystems, #AM1560) following the manufacturer’s protocol. Total RNA was used to construct sequencing libraries. mRNA was enriched using the NEBNext Poly(A) mRNA Magnetic Isolation Module (E7490S), and libraries were prepared with the NEBNext UltraExpress RNA Library Prep Kit (E3330S). Sequencing was performed on an Illumina NovaSeq 6000 system to generate 150-base-pair paired-end reads.

### Native-ChIP-seq

Native ChIP-seq was carried out as previously described^56^. Cells were lysed in 0.1% Triton X-100 and 0.1% Sodium Deoxycholate supplemented with a protease inhibitor cocktail. Chromatin in the lysate was digested with micrococcal nuclease (MNase, New England BioLabs, #M0247S) at room temperature for 5 minutes, and the reaction was stopped with 5 µl of 0.25 mM EDTA. Preclearing of the digested chromatin was done for 2 hours using anti-IgA magnetic beads (Dynabeads, ThermoFisher, #10001D), followed by overnight incubation with antibody-bead complexes containing H3K27ac antibody (Diagenode, #C15410196) in immunoprecipitation (IP) buffer (20 mM Tris-HCl pH 7.5, 2 mM EDTA, 150 mM NaCl, 0.1% Triton X-100, 0.1% sodium deoxycholate) at 4 °C.

IPs were sequentially washed twice with low-salt buffer (20 mM Tris-HCl pH 8.0, 2 mM EDTA, 150 mM NaCl, 1% Triton X-100, 0.1% SDS) and high-salt buffer (20 mM Tris-HCl pH 8.0, 2 mM EDTA, 500 mM NaCl, 1% Triton X-100, 0.1% SDS). Complexes were eluted in elution buffer (1% SDS, 100 mM sodium bicarbonate) for 1.5 hours at 65°C. Histones were digested with protease (Qiagen, #19155) for 30 minutes at 50 °C, and DNA fragments were purified using Sera-Mag magnetic beads in 30% PEG. Illumina libraries were prepared as previously described by end repair, 3′ A-tailing, and adaptor ligation (New England BioLabs, #E6000B-10). Libraries were indexed and amplified by PCR (10 cycles) before sequencing on the Illumina HiSeq 2500 or NovaSeq 6000 according to the manufacturer’s instructions.

### CUT&RUN-seq

CUT&RUN was performed using the EpiCypher CUTANA kit (14-1048) according to the manufacturer’s protocol with the following modifications^57^. Freshly sorted epithelial cells were resuspended in nuclei isolation buffer (20 mM HEPES, 20 mM KCl, 0.5 mM spermidine, 0.05% IGEPAL, 50% glycerol, 1X protease inhibitor in ddH2O). Cells were washed twice in this buffer and centrifuged at 800g for 5 minutes. The isolated nuclei were resuspended in permeabilization buffer (CUTANA Wash Buffer, 0.01% digitonin, 1X protease inhibitor), incubated on ice for 5 minutes, and captured on Concanavalin A beads at room temperature for 10 minutes. Captured nuclei were then resuspended in 180 µl of antibody buffer (permeabilization buffer with 2 mM EDTA) containing 0.5 µg of Sox17 antibody (RnD, #AF1924) or β-catenin antibody (Thermo Fisher, #71-2700) and incubated overnight. For Sox17 CUT&RUN, a secondary antibody, Rabbit Anti-Goat IgG H&L (Abcam, #ab6697), was added for 2 hours the following day. IgG control reactions were performed in parallel for each genotype and cell population by omitting the primary antibody and incubating nuclei with Rabbit Anti-Goat IgG H&L (Abcam, #ab6697) alone, otherwise following the identical protocol, library construction, sequencing and normalisation. Antibody-bound chromatin was digested at 4 °C for 30 minutes on a nutator, followed by fragment release for an additional 30 minutes at 4 °C. DNA fragments were purified using SeraMag beads with 30% PEG-8000. Library construction was performed as described in the Native ChIP-seq section.

### ATAC-seq

Accessible chromatin was profiled essentially as described by Buenrostro and colleagues^58^. For each replicate, 35,000 to 50,000 viable cells were collected, rinsed twice in ice-cold PBS and pelleted at 500 × g for 5 min at 4 °C. Pellets were taken up in 50 μL of cold buffer containing 10 mM Tris-HCl pH 7.4, 10 mM NaCl, 3 mM MgCl₂ and 0.1% (v/v) IGEPAL CA-630, and nuclei were recovered by a second spin at 500 × g for 10 min at 4 °C. Nuclei were then transposed directly in a 50 μL reaction with Tn5 (Diagenode #C01070012) for 30 min at 37 °C with gentle agitation, and the tagmented DNA was recovered on a silica column (Zymo). Libraries were generated from the purified DNA using a shared forward primer and sample-specific barcoded reverse primers in a high-fidelity PCR master mix (New England Biolabs). Reactions were first taken through five cycles; a 5 μL aliquot was then monitored by qPCR to establish the cycle number corresponding to one third of maximum fluorescence, and the remaining material was amplified for that number of additional cycles. Products were column purified, and fragments below 100 bp were depleted with AMPure XP beads (Beckman Coulter). Library concentration was measured with the Qubit dsDNA High Sensitivity assay (Thermo Fisher) and fragment-size distribution was confirmed before pooling. Sequencing was carried out on an Illumina NovaSeq 6000 platform.

### Co-immunoprecipitation and western blotting

Epithelial cells isolated from Apc^−/−^ adenomas were lysed for 1 h at 4 °C in lysis buffer (10 mM HEPES-KOH pH 7.9, 1.5 mM MgCl₂, 10 mM KCl, 1 μM dexamethasone, 0.5 mM DTT, 0.15% NP-40, protease inhibitors (Calbiochem #539134), passed through a 40 μm filter and centrifuged at 21,000 × g for 10 min at 4 °C. Whole-cell lysates were precleared for 2 h with 20 μL anti-IgG/IgA magnetic beads (Dynabeads, Thermo Fisher #10001D), then incubated overnight at 4 °C with 2 μg anti-SOX17 (R&D #AF1924) or rabbit anti-goat IgG H&L (Abcam #ab6697) and 20 μL beads in IP buffer (75 mM KCl, 20 mM HEPES, 2.5 mM MgCl₂, 1 mM DTT, 0.1% NP-40). Immuno-precipitates were washed five times in IP buffer, eluted in 1× NuPAGE LDS Sample Buffer (Thermo Fisher #NP0007) and heated at 95 °C for 5 min. Input corresponds to 10% of the lysate used per immunoprecipitation.

For western blotting, 30 μL of each eluate was resolved on Mini-PROTEAN TGX gels (Bio-Rad #4569033) for 1 h on ice and transferred with a Trans-Blot Turbo system (Bio-Rad #1704150EDU). Membranes were probed overnight at 4 °C with anti-β-catenin (Thermo Fisher, #71-2700), washed five times in TBS-T (20 mM Tris pH 7.5, 150 mM NaCl, 0.1% Tween 20), incubated for 1 h at room temperature with anti-rabbit IRDye 680RD (LI-COR #926-68071) and anti-goat IRDye 800CW (LI-COR #926-32210), and imaged on a LI-COR Odyssey CLx.

All cartoon illustrations in figures were created with BioRender.com.

## QUANTIFICATION AND STATISTICAL ANALYSIS

All bioinformatics analyses were conducted on R (v.4.1.0).

### Single cell multiome preprocessing, alignment and filtering

Single-cell multiome reads were aligned to the mouse mm10 genome using Cell Ranger Arc (10X Genomics, v2.0.0). Analysis and stringent quality control of scRNA-seq and scATAC-seq data were performed using Seurat v4^15^ and Signac v1.3.0^16^, respectively. For scRNA-seq, cells were retained if they met the following criteria: >12,000 unique molecular identifiers (UMIs) per cell, >1,000 genes detected, mitochondrial transcript ratio <0.35 per cell, and expression novelty >0.8. scATAC-seq filtering criteria included cells with >40% reads in peaks, >1,000 fragments per cell, blacklist fraction <0.05, transcription start site (TSS) enrichment >2, and nucleosome signal <4. After filtering, 8,419 cells were included for analysis, comprising 4,511 *WT* cells and 3,908 day 14 *Apc^−/−^* cells. ATAC peaks were called using the ‘CallPeaks’ function in MACS2 via Signac, and nucleosome signal and TSS enrichment were calculated using the ‘NucleosomeSignal’ and ‘TSSEnrichment’ functions.

Filtered scRNA-seq data were normalized using SCTransform (SCT) followed by principal component analysis (PCA). For scATAC-seq, latent semantic indexing (LSI) was applied to the top variable peaks, combining term frequency–inverse document frequency normalization with singular value decomposition for dimensionality reduction. The first LSI component was excluded due to correlation with sequencing depth. PCA components from scRNA-seq and LSI components from scATAC-seq were integrated in ‘FindMultiModalNeighbors’ to construct a joint neighbor graph. A multimodal weighted k-nearest neighbor algorithm was then applied to generate a Uniform Manifold Approximation and Projection (UMAP) embedding using ‘RunUMAP’. Finally, cell clusters were identified from the combined UMAP using ‘FindClusters’.

### Single cell multiome cell type annotation

Top differential markers for each identified cluster or sub-cluster were determined using the ‘FindMarkers’ function with an adjusted p-value <0.05, minimum 1.2-fold change, and expression in ≥10% of cells within the comparison groups. The Wilcoxon rank-sum test was used, followed by Bonferroni correction for multiple comparisons. Cell identities were assigned based on multiple cell type-specific or enriched marker genes previously described in the literature^59^. These included but were not limited to *Lgr5*, *Smoc2, Cdca7, Olfm4, Ascl2, Soat1* for Stem cells, *Mki67, Ube2c, Top2a, Pcna, Mcm6, Cdk4, Mcm5, Ccna2, Cdk1* for Transit amplifying cells*, Lyz1, Defa17, Defa22, Ang4* for Paneth cells, *Chga, Cck,Ghrl, Neurod1, Chgb* for enteroendocrine cells, *Muc2, Spdef, Clca3, Agr2* for goblet cells, *Alpi, Fabp6, Apoa1, Apoa4* for enterocytes.

### Differentially accessible regions

Differential peaks for each cluster in the scATAC-seq were identified using logistic regression model in ‘FindMarkers’ function with *padj* < 0.05, logFC > 0.25, min pct >10% and using “nCount_peaks” which is the total number of fragment counts per cell for variables to test.

### Gene expression and Chromatin accessibility score

Gene expression signature scores for specific gene sets were calculated using the ‘AddModuleScore’ function in Seurat, and cells were annotated based on their respective maximum module score. This score reflects single-cell expression levels for each cluster, normalized by the aggregated expression of control feature sets. Features were binned according to average expression, and control features were randomly selected from each bin. For chromatin accessibility, cluster-specific differential peak sets were scored using the ‘AddChromatinModule’ function in Signac, generating a combined accessibility score that was incorporated into the Seurat object.

### Functional enrichment of differentially expressed genes (DEGs)

Enrichment analysis of DEGs was performed using the Metascape web tool (www.metascape.org). The gene sets obtained from the GO processes, KEGG pathways, Reactome gene sets, canonical pathways, and CORUM complexes were used. *P*-values were calculated based on the accumulative hypergeometric distribution and Benjamini-Hochberg *p*-value correction algorithm. Terms with *p*-value < 0.01 were collected and grouped into clusters based on their membership similarities.

### scATAC–seq motif analysis

Motif position frequency matrices from JASPAR 2020 were added to the data using Signac. ChromVAR motif analysis was performed using ‘RunChromVAR’ in Signac to calculate per cell motif activity. The results of selected TFs were visualized on UMAP using ‘FeaturePlot’.

### Peak-to-gene linkage analysis

For each peak–gene pair, a linkage score was calculated in Signac using the ‘LinkPeaks’ function. The Pearson correlation coefficient (r) was computed between gene expression and the accessibility of each peak located within 500 kb of the gene TSS. A background set of expected correlations was generated by randomly sampling 200 peaks on different chromosomes from the gene, matched for GC content, accessibility, and sequence length using the MatchRegionStats function in Signac. *P*-values for each peak were determined using a one-sided z-test. Peak–gene links with p-values <0.05 and a Pearson correlation coefficient >0.05 or <−0.05 were retained and visualized using the CoveragePlot function in Signac.

### Single cell pseudotime analysis

Trajectory analysis was performed using the Monocle3 (v.1.4.26) using both scRNA-seq and scATAC-seq data. Seurat object was converted to CellDataSet used by Monocle3 using conversion function in SeuratWrappers (v.0.4.0). UMAPs for scRNA and scATAC were used in ‘cluster_cells’. ISCs or *Ad-Stem* were designated as root cells for pseudotime estimates for *WT* and *Apc^−/−^*samples respectively. Pseudotime values were extracted and added to the Seurat object and visualized using Seurat functions. For the binned comparison in Extended Data Fig. 2e–g, wild-type stem, transit-amplifying and enterocyte cells were ordered along a transcriptional trajectory using Monocle 2 (v.2.36.0), with DDRTree applied to the 500 most variable genes and the stem population as root, and divided into five bins of equal cell number. Enterocyte signature scores were computed with Seurat AddModuleScore from nine enterocyte genes (*Alpi*, *Apoa1*, *Apoa4*, *Fabp1*, *Rbp2*, *Lct*, *Slc5a1*, *Fabp2*, *Ada*), applied to the RNA assay and to the peaks assigned to those genes with ClosestFeature; a cell was scored as having reached enterocyte level if its score exceeded the 25th percentile of mature enterocytes for that modality.

### mRNA-seq

Raw RNA-seq reads were aligned to the mouse genome (GRCm38, GENCODEvM25) using STAR aligner v2.7.8a in 2-pass mode. Quality assessment included per-base sequence quality, per-read GC content (∼50%), strand-specific alignment distribution, exon versus intron read distribution, and 3′ bias. Transcript abundance was quantified as read counts using HTSeq v0.13.5 and normalized across libraries with DESeq2, then converted to reads per kilobase of transcript per million mapped reads (RPKM). Differential expression between samples was determined using DESeq2, applying false-discovery rate (FDR) thresholds as indicated. All alignment and initial processing steps were executed using the Snakemake workflow management system v6.6.1. Visualization of gene expression data was performed using ggplot2 v3.5.2 and pheatmap v1.0.13.

### ChIP-seq

ChIP-seq data were trimmed using trim-galore (v0.6.6) to remove adapters and low-quality reads. The cleaned reads were aligned to the reference genome using bwa-mem v0.7.17 with default settings. Reads that were non-uniquely mapped, duplicates, or had a MAPQ score below 5 were removed using Samtools v1.17. Peaks for H3K27ac were called with MACS v2.2.9.1 in sharp mode using a q-value cutoff of 0.01. BigWig files were generated with deepTools v3.5.5 using a bin size of 10, smooth length of 10, and quantile normalization via Haystack. Blacklisted regions defined by the ENCODE consortium were removed using Bedtools v2.30.0.

### Motif enrichment analysis for CUT&RUN and ChIP-seq

Motif analysis was performed using findMotifsGenome.pl from HOMER (v5.1) on the peak summit (-size 200).

### Determination of differential genomic binding

Peaks for H3K27ac ChIP-seq and for SOX17 and β-catenin CUT&RUN were called with MACS v2.2.9.1 in narrow-peak mode using a q-value cutoff of 0.01. Blacklisted regions defined by the ENCODE consortium were excluded before any comparison. For differential analyses, signal was quantile normalised across samples using the Haystack algorithm^60^. Differential signal between conditions was then determined on the quantile-normalised counts within the union peak set using a Wald test in DESeq2. Regions were partitioned into the classes shown in each figure by k-means clustering of the normalised signal across the samples being compared. For the β-catenin CUT&RUN comparison across WT intestinal stem cells and day 14 *Apc ^−/−^* stem and non-stem cells (Fig. 2b), clustering was performed across all three cell types and classes were required to show a significant change in signal at *q < 0.01*. For the stratification of β-catenin binding into SOX17-dependent and SOX17-independent classes (Fig. 3f), regions were required to show at least a two-fold change in signal between any two cell types. For the H3K27ac enhancer clusters across *WT*, *Apc ^−/−^* and *Apc ^−/−^Sox17 ^−/−^* stem cells (Fig. 4a), regions were required to show at least a two-fold change in signal at *q < 0.05*. The number of regions in each resulting class is given in the corresponding figure legend and in Results. Comparisons of target signal against IgG were performed on 1× genome-coverage normalised tracks without quantile normalisation, since quantile normalisation equalises the signal distributions of the samples being compared and would artificially raise the IgG control toward its target. Stratification of differentially accessible regions in the single-cell ATAC data (Fig. 1e) is described separately under “Differentially accessible regions”.

### TCGA bulk RNA-seq data processing and CMS classification

Raw gene counts for TCGA-COAD were obtained and aligned to the human genome (GRCh38, GENCODEv38) using STAR aligner v2.7.8a. Raw counts from the TCGA-COAD dataset were used as input for CMScaller^25^, and CMS classification was performed with default parameters. CMS-annotated patient samples were then analyzed with DESeq2 to perform differential expression analysis, generate normalized counts, and convert values to RPKM.

### Data processing of scRNA-seq human dataset

Raw count matrices for the SMC samples^29^ were retrieved from the Gene Expression Omnibus (GEO, accession no. GSE132465). Seurat objects were generated from these matrices, and the dataset was filtered to include only epithelial cells based on available annotations. Data were processed and analyzed following the same workflow as described for the mouse scRNA-seq. Metadata, including cell type annotations and CMS classifications, were obtained from the original publication.

### Statistics and reproducibility

Experiments were reproduced as described in the figure legends, with representative panels repeated at least two to three times. Animal numbers and are specified in the respective figure legends. Statistical analyses were performed using R (v4.1.0). Computational figures were generated with ggplot2 (v3.5.2), ggpubr (v0.6.2), patchwork (v1.3.2), Seurat (v4.4.0), and Signac (v1.16.0).

All statistical tests are indicated in the figure legends. Significance levels are defined as follows: NS (not significant): *P*> 0.05, ^∗^*P* ≤ 0.05, ^∗∗^*P* ≤ 0.01, and ^∗∗∗^*P* ≤ 0.001.

## Acknowledgments

This work was supported by NIH grants K01DK113067, R03DK134799, Donald E. and Delia B. Baxter Foundation Fellowship Award, Tower Cancer Research Foundation Fellowship Award, and The Uri and Karen Geiger Cancer Metastasis Research and Innovation Fund to U.J. A.L was supported by Canadian Institute of Health Research (CIHR) Post-Doctoral Fellowship and California Institute of regenerative Medicine (CIRM) Post-Doctoral Fellowship. A.L was supported by Canadian Institute of Health Research (CIHR) Post-Doctoral Fellowship, California Institute of regenerative Medicine (CIRM) Post-Doctoral Fellowship and NCI grant K99CA293236.

## Contributions

S.S. and U.J. conceptualized the study design. S.S., A.L., M.B., and E.S. performed experimental work. S.S. and G.Y. performed the computational and data analysis. S.S. and J.V. performed image analysis and quantification. S.S. and U.J. wrote the manuscript. All authors read and approved the final manuscript.

---

**Extended Data Figure 1.**
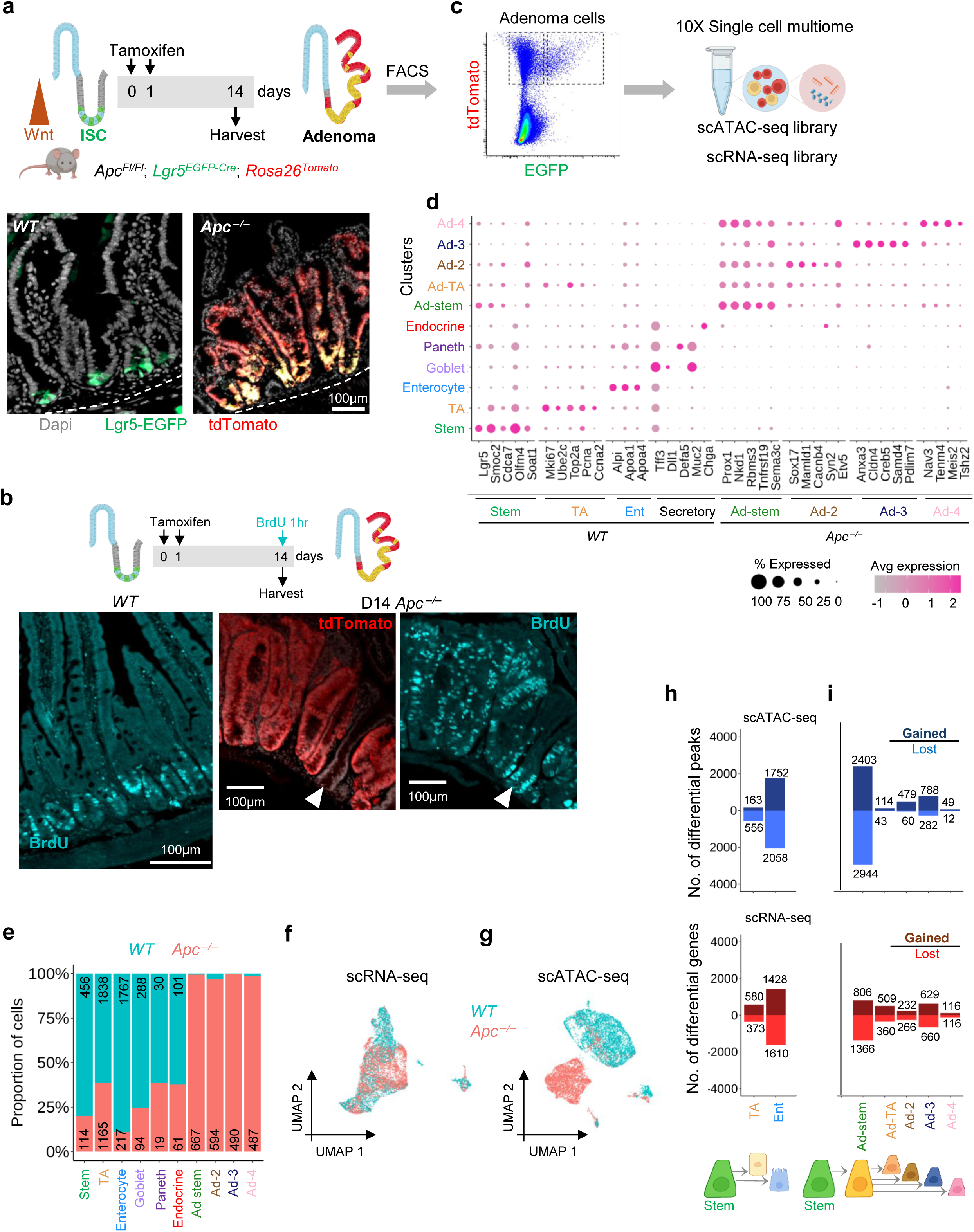
**a.** Experimental schematic illustrating two consecutive intraperitoneal Tamoxifen (TAM) injections that induce *Apc* deletion in Lgr5^+^ ISCs (green), driving adenoma formation throughout the intestine and labeling all transformed cells with tdTomato (red). Tissues are collected 14 days after TAM administration. Immunofluorescence images below demonstrate expansion of Lgr5⁺ ISCs and lineage tracing of transformed daughter cells marked by tdTomato in *WT* and *Apc^−/−^*mouse small intestines. Dotted lines indicate muscle layers. **b.** Immunohistochemistry (IHC) with fluorescent staining shows increased proliferative cells in *Apc^−/−^* mouse intestine 14 post-TAM based induction of adenomas, compared to the crypt-restricted proliferation in *WT* marked by BrdU (injected 1 hr prior to harvest). White arrows in D14 *Apc^−/−^*intestine indicate normal crypts with limited spread of BrdU^+^ proliferating cells. **c.** Experimental workflow illustrating isolation of adenoma cells and 10x scMultiome pipeline based library generation. **d.** Dot plot showing expression of known marker genes in cell clusters identified in *WT* and *Apc^−/−^* cells. Dot size represents percentage of cells expressing the gene. **e.** Bar plot showing changes in proportion of different cell types between *WT* and *Apc^−/−^* samples based on scMultiome data. Numbers within bars represent total cells in each cluster. **f-g.** UMAPs for *WT* and *Apc^−/−^* cells generated using only scRNA-seq **(f)** or scATAC-seq **(g)** features. **h-i.** Barplot showing the number of differential ATAC peaks (*padj* <0.05, 1.2-fold change, logistic regression test) (Top) and genes (*padj* <0.05, 1.2-fold change, Wilcoxon Rank Sum test) (Bottom) that are gained or lost during ISC differentiation into TA or enterocyte cells (**h**) and during ISC transformation upon *Apc* deletion and further into various transformed cell cultures in adenomas (**i**).

**Extended Data Figure 2.**
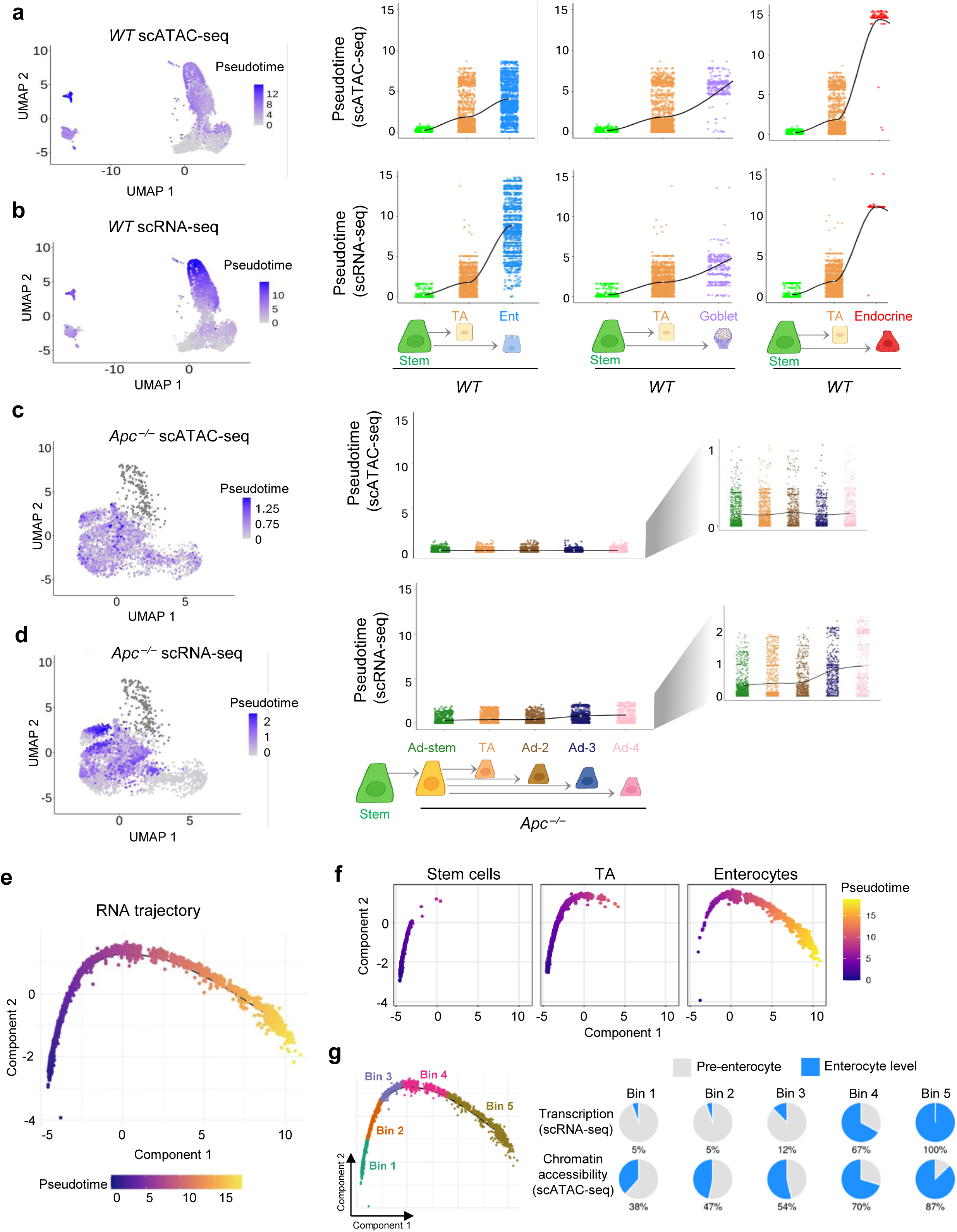
a-b. UMAP plots showing pseudotime projections (Monocle 3) of *WT* cells based on scATAC-seq (**a**) or scRNA-seq (**b**), inferring differentiation trajectories from stem cells to differentiated states. Plots on the right show Monocle 3-derived cell differentiation trajectories for *WT* cell types (clusters), with increasing pseudotime along the scATAC-seq (**a**) and scRNA-seq (**b**) axes. ISC differentiation into absorptive (enterocytes) and secretory (goblet, and endocrine) lineages follows the expected progression along. **c-d.** UMAPs showing pseudotime projections of *Apc ^−/−^* adenoma cells based on scATAC-seq (**c**) or scRNA-seq (**d**) reveal a lack of detectable differentiation trajectories or pseudotime progression compared to the *WT* cells (**a-b**). Similarly, transformed stem (*Ad-stem*) cells show no appreciable pseudotime increase toward other adenoma clusters (*Ad-2, Ad-3, Ad-4*), suggesting that independent, parallel cell trajectories may underlie adenoma cell heterogeneity. **e–g.** Wild-type cells ordered by RNA pseudotime (Monocle 2). **e.** Transcriptional trajectory coloured by pseudotime. **f.** The same trajectory showing stem, transit-amplifying and enterocyte cells separately. **g.** Cells divided into five bins of equal cell number (n = 812–813 per bin), shown along the trajectory (left) and, for each bin, the fraction of cells that have reached enterocyte-level signal (right). Scores were computed from a single set of enterocyte genes (*Alpi*, *Apoa1*, *Apoa4*, *Fabp1*, *Rbp2*, *Lct*, *Slc5a1*, *Fabp2*, *Ada*), either as chromatin accessibility at the associated peaks or as expression of those genes, and a cell was counted as enterocyte-level if its score exceeded the 25th percentile of mature enterocytes for that modality. Enterocyte-level accessibility is present in 38% of cells in Bin 1, where 5% have reached enterocyte-level expression.

**Extended Data Figure 3.**
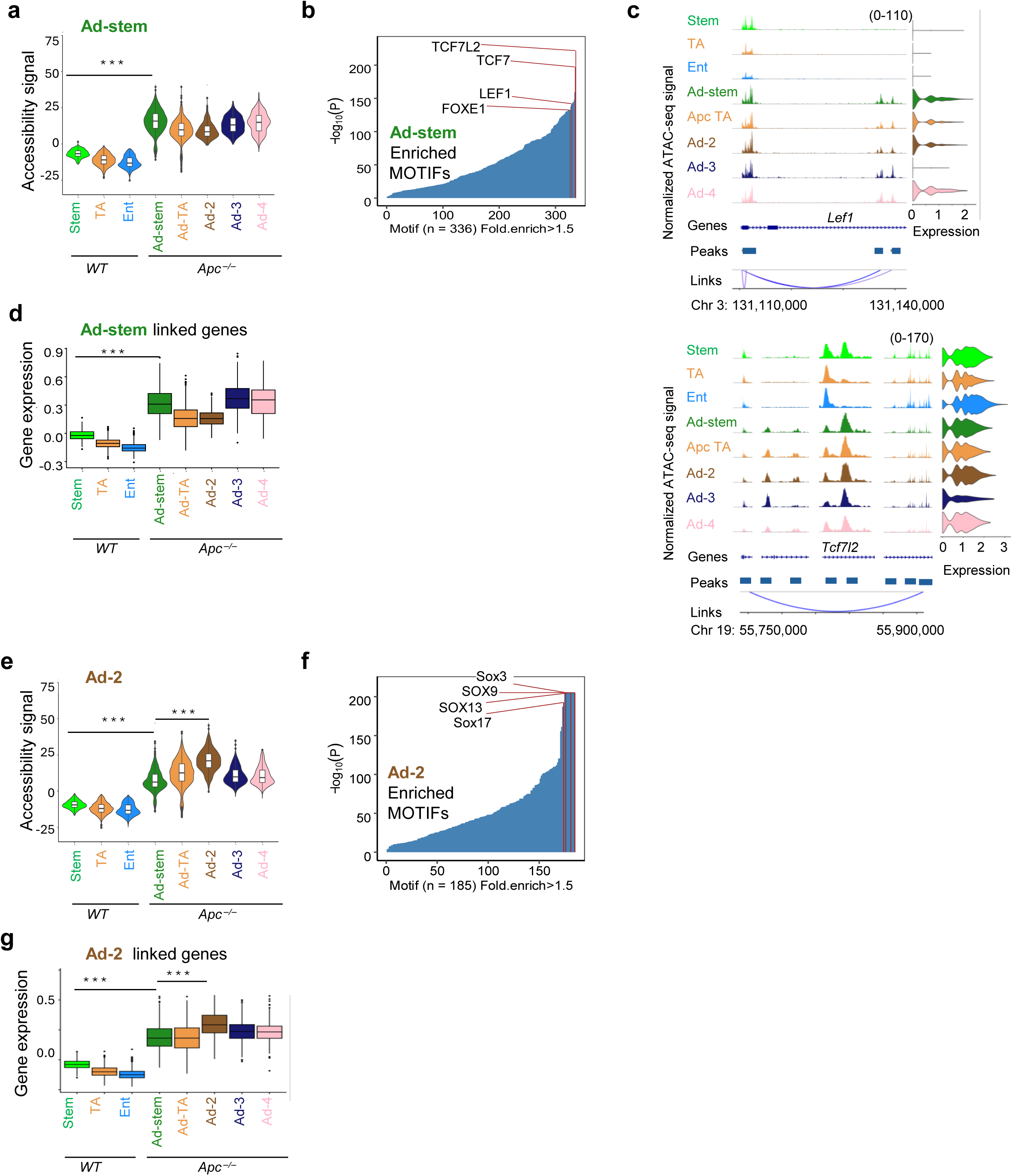
**a.** Violin plots showing average scATAC-seq accessibility signal for differential peaks specific to *Ad-stem* cluster; Wilcoxon test *p*-value is reported. (*** *p* < 0.001). **b.** Motif enrichment plot show distinct TF motifs enriched in *Ad*-stem cluster with ≥1.5 or higher fold enrichment. The number of Motifs plotted for the cluster are indicated. Motifs are ranked by significance (*p* value from a hypergeometric test). **c.** Coverage plots show normalized scATAC-seq signal for each cluster across genomic loci of *Lef1* (Top) and *Tcf7l2* (Bottom). Blue bars mark scATAC-called peaks, and gradient blue links indicate correlations between peaks and their associated genes. Violin plots on the right display gene expression for each locus. **d.** Boxplot showing comparative expression levels (average expression score across cells within the indicated clusters) of genes linked to enhancers specifically accessible in *Ad-stem* cells. Wilcoxon test *p*-value is shown. (*p* < 0.001 = ***). **e.** Violin plots showing average scATAC-seq accessibility signal for differential peaks specific to *Ad-2* cluster; Wilcoxon test *p*-value is reported. (*** *p* < 0.001). **f.** Motif enrichment plot show distinct TF motifs enriched in *Ad-2* cluster with ≥1.5 or higher fold enrichment. The number of Motifs plotted for the cluster are indicated. Motifs are ranked by significance (*p* value from a hypergeometric test). **g.** Boxplot showing comparative expression levels (average expression across cells within the indicated clusters) of genes linked to enhancers specifically accessible in *Ad-2* cells. Wilcoxon test *p*-value is shown. (*p* < 0.001 = ***).

**Extended Data Figure 4.**
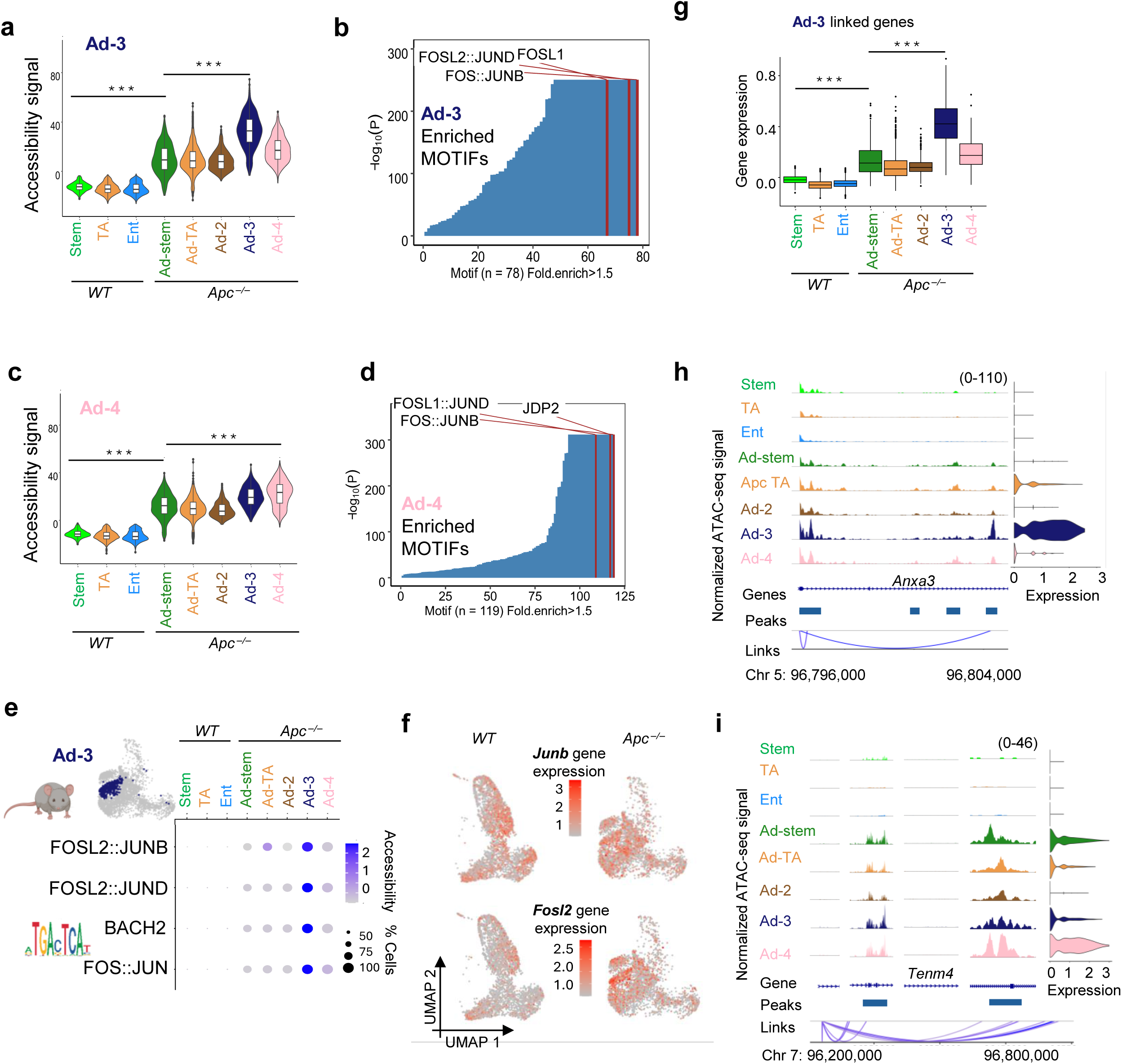
**a.** Violin plots showing average scATAC-seq accessibility signal for differential peaks specific to *Ad-3* cluster; Wilcoxon test *p*-value is reported. (*** *p* < 0.001). **b.** Motif enrichment plots show distinct TF motifs enriched in *Ad-stem* cluster with ≥1.5 or higher fold enrichment. The number of Motifs plotted for the cluster are indicated. Motifs are ranked by significance (*p* value from a hypergeometric test). **c.** Violin plots showing average scATAC-seq accessibility signal for differential peaks specific to *Ad-4* cluster; Wilcoxon test *p*-value is reported. (*** *p* < 0.001). **d.** Motif enrichment plots show distinct TF motifs enriched in *Ad-stem* cluster with ≥1.5 or higher fold enrichment. The number of Motifs plotted for the cluster are indicated. Motifs are ranked by significance (*p* value from a hypergeometric test). **e.** Dot plot showing significant gain of accessibility at loci with JUNJ/FOS motifs in *Apc ^−/−^* cells, particularly *Ad-3* cells, compared to *WT* cells. Dot size represents the percentage of cells enriched for the TF motifs in *WT* and *Apc ^−/−^* cell clusters. **f.** Split UMAP of *WT* and *Apc ^−/−^* cells showing expression of *Junb* (Top) and *Fosl2* (Bottom). Heat indicates gene expression level. **g.** Boxplot showing comparative expression levels (average expression score across cells within the indicated clusters) of genes linked to enhancers specifically accessible in *Ad-3* cells. Wilcoxon test *p*-value is shown. (*** *p* < 0.001). **h-i.** Coverage plots show normalized scATAC-seq signal for each cluster across genomic loci of *Anxa3* (**h**) and *Tenm4* (**i**). Blue bars mark scATAC-called peaks, and gradient blue links indicate correlations between peaks and their associated genes. Violin plots on the right display gene expression for each locus.

**Extended Data Figure 5.**
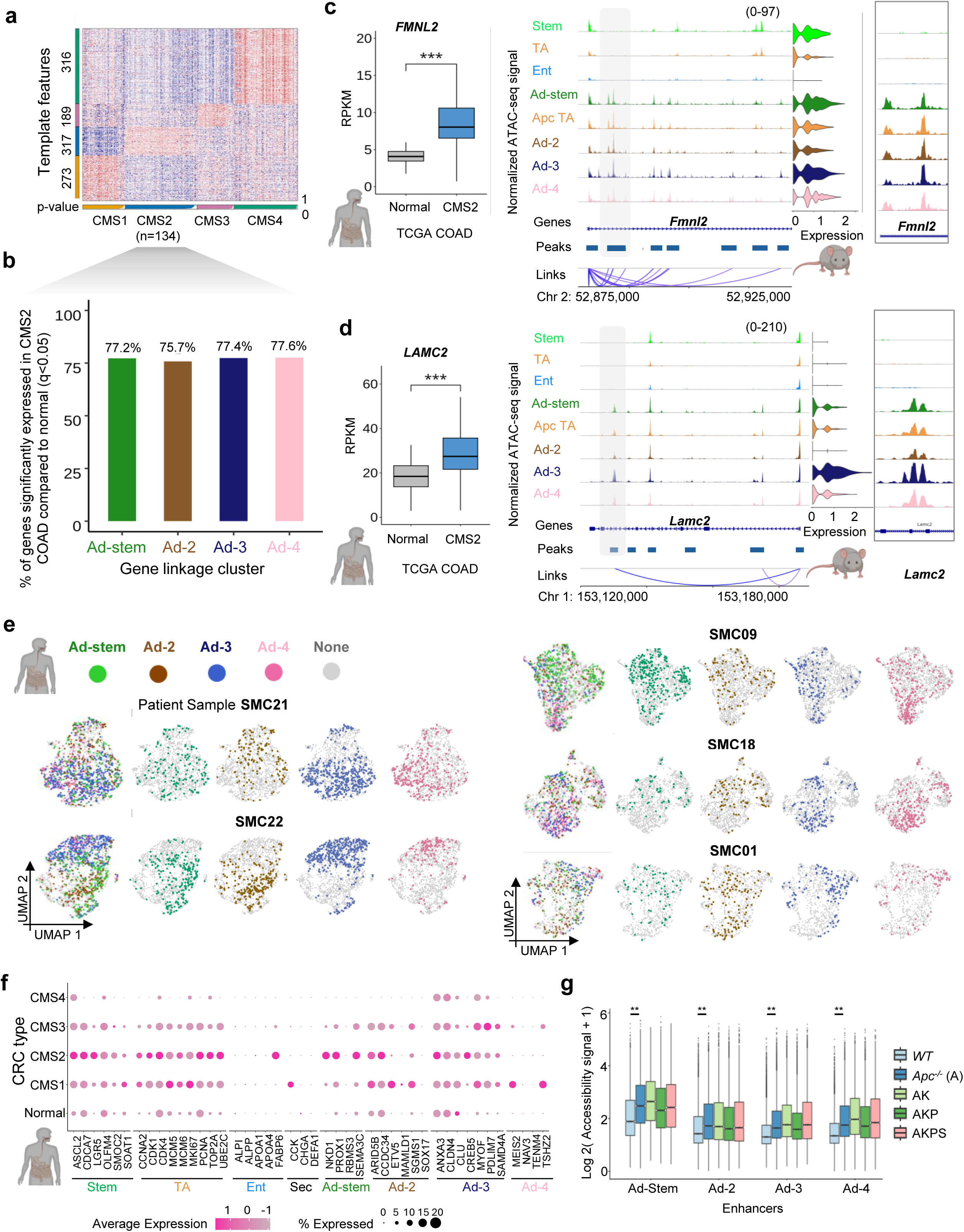
**a.** Heatmap showing CMS subtypes of human CRC tumour samples (TCGA-COAD mRNA-seq data) as determined using the CMS caller algorithm, which uses genes indicated on the Y-axis for CMS classification. Subtypes are highlighted in yellow (CMS1), blue (CMS2), pink (CMS3), and green (CMS4). The *p-*value scale ranges from 0 to 1. **b.** Bar plot showing percentage of genes specifically activated in adenoma clusters of *Apc ^−/−^* mice that exhibit significantly higher expression in CMS2 tumours compared to normal samples in TCGA-COAD (*q < 0.05,* DEseq2). **c.** Boxplot (left) showing *FMNL2* expression in TCGA-COAD CMS2 tumours versus normal samples (t-test, *p < 2.5e-23*) Coverage plot (middle) highlights normalized ATAC-seq signal across clusters for *Fmnl2* in *WT* and *Apc ^−/−^* scMultiome data. Tracks on the right show a zoomed-in view of the enhancer highlighted in grey. **d.** Boxplot (left) showing gene expression of *LAMC2* in TCGA-COAD CMS2 tumours versus normal samples (t-test, *p < 9.5e-07*). Coverage plot (middle) highlights normalized ATAC-seq signal across clusters for *Lamc2* in *WT* and *Apc ^−/−^* scMultiome data. Tracks on the right show a zoomed-in view of the enhancer highlighted in grey. **e.** UMAP plots for individual patient samples (containing substantial numbers of CMS2 epithelial cells) from *Lee et al.* 2020^29^ showing expression of genes linked to adenoma clusters (*Ad-stem, Ad-2, Ad-3,* and *Ad-4*). Cells with the average gene scores > 0.05 were considered positive for the signature. **f.** Dot plot showing expression of marker genes for each cluster from Extended Data Fig. 1d and top genes expressed in adenoma cell clusters in epithelial cells from normal human colon or from various CRC subtypes. Patient samples are from *Lee et al.* 2020^29^. Dot size represents the percentage of cells expressing the gene. **g.** Boxplots quantifying bulk ATAC-seq signal (log2(mean matrix signal + 1)) at the Ad-stem, Ad-2, Ad-3 and Ad-4 enhancer sets (defined in Fig. 1e) across *WT*, *Apc^−/−^* (A), *Apc^−/−^*;*Kras^G12D^* (AK), *Apc^−/−^*;*Kras^G12D^*;*Trp53*^−/−^ (AKP) and *Apc^−/−^*;*Kras^G12D^*;*Trp53*^−/−^;*Smad4^−/−^*(AKPS) intestinal stem cell organoids (GSE241382). Signal was averaged across independent organoid cultures per genotype (n = 4 WT, 3 A, 4 AK, 2 AKP, 3 AKPS) and each point is one enhancer (n = 4,344 Ad-stem, 5,873 Ad-2, 10,044 Ad-3, 5,658 Ad-4). Boxes show the median and interquartile range; whiskers extend to 1.5 × IQR. Brackets show the *WT* to *Apc^−/−^*comparison within each enhancer set, two-sided Wilcoxon rank-sum test across enhancers with Benjamini–Hochberg correction; ** denotes *p* < 2.2 × 10^−16^, the value obtained for all four sets.

**Extended Data Figure 6.**
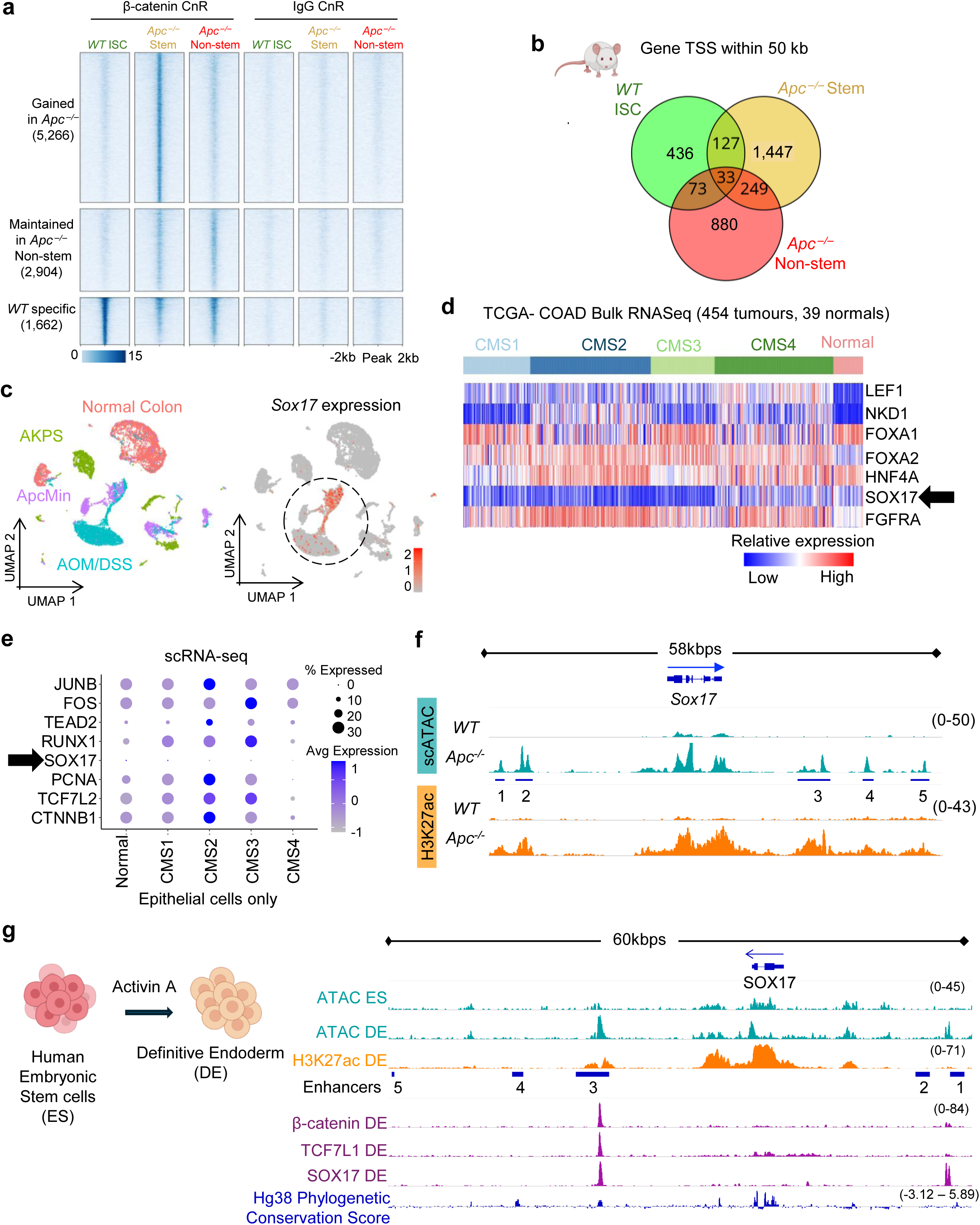
**a.** Heatmap showing β-catenin CUT&RUN signal at the peak classes in Fig. 2b, with the matched IgG CUT&RUN control from the same genotypes and cell populations alongside. IgG signal is flat across all classes. **b.** Venn diagram showing the overlap of genes located within 50 kb of the β-catenin bound enhancers across different cell types in mouse tissues. **c.** UMAP (left) showing mouse adenoma scRNA-seq data from *Tian et al.,* 2023^41^; *Sox17* expression is shown in red in the UMAP on the right. The dotted circled area highlights early adenoma cells from different models showing Sox17 expression. **d.** Heatmap showing expression of various genes in TCGA-COAD tumours classified into CMS subtypes and in normal samples. **e.** Dot plot showing scRNA-seq expression of genes in epithelial-only CRC patient cells from *Lee, et al., 2020*^29^. Heat represents average expression level, and dot size represents percentage of cells expressing the gene. **f.** Genome browser view of 58 kb window around the *Sox17* gene in the mouse genome showing scATAC-seq peaks and H3K27ac signal in *WT* ISCs and *Apc ^−/−^* stem cells. Enhancers activated in *Apc ^−/−^*stem cells are numbered and highlighted. **g.** Schematic of hESCs differentiation to DE samples used for the analysis of data from *Mukherjee et al.,* 2022^39^ and UCSC genome browser. Genomic tracks show enhancers around human Sox17 gene (corresponding to mouse enhancers in panel (**f**), identified using LiftOver). Tracks display ATAC-seq, H3K27ac, β-catenin, TCF7L1 and SOX17 signals in DE across the loci. Vertebrate conservation scores are shown in blue.

**Extended Data Figure 7.**
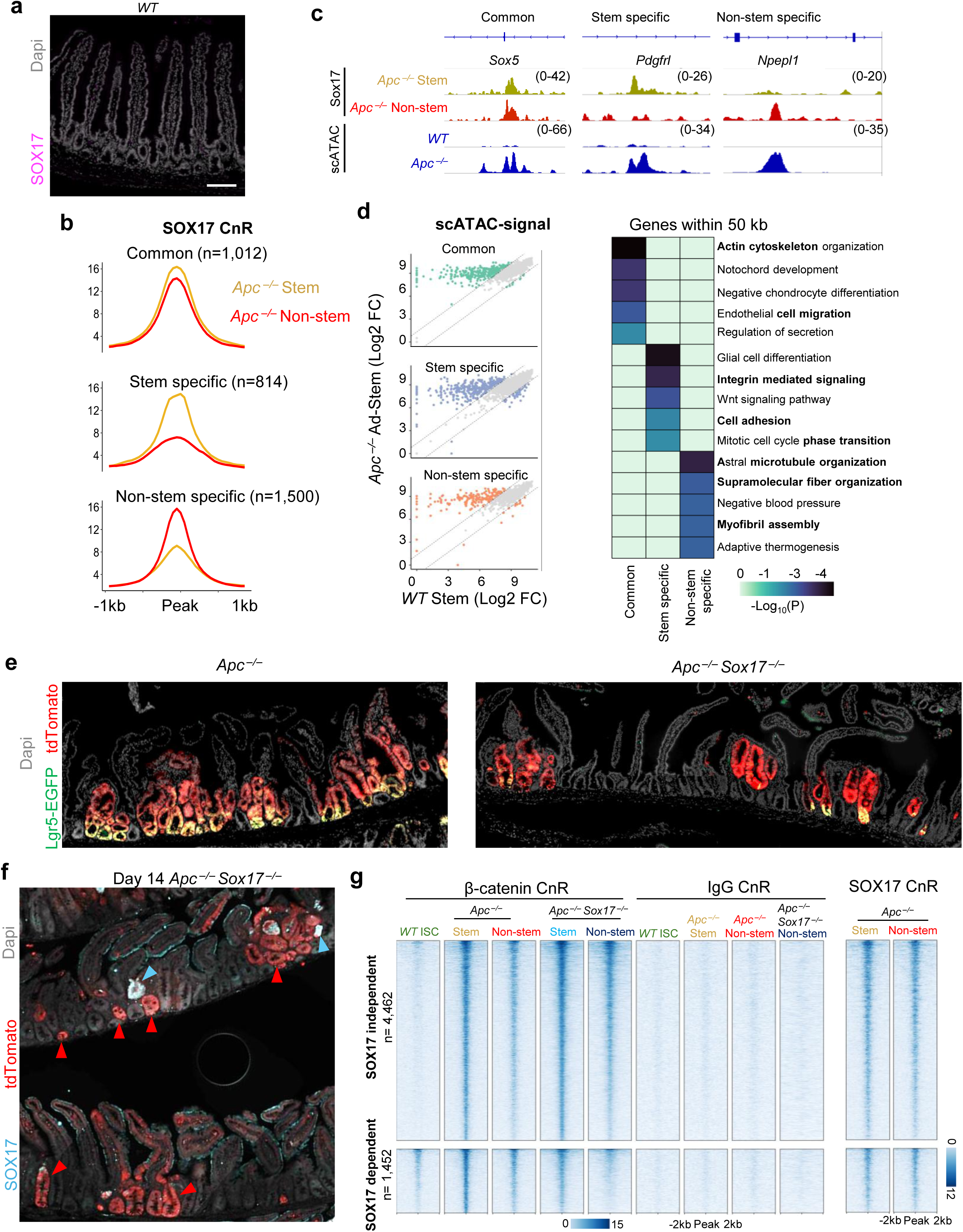
**a.** Fluorescent image showing immunohistochemical staining of SOX17from wild-type mouse intestine, demonstrating that Sox17 is not expressed in normal intestinal epithelial cells. Scale bar = 100 μm. **b.** Profile plot showing SOX17 CUT&RUN-seq (CnR) signal in *Apc ^−/−^*stem and non-stem cells for enhancers with increased or reduced binding across cell types (multibigwig summary of all peaks). **c.** Genome browser traces showing examples of SOX17-bound CnR peaks, scATAC-seq signal, and H3K27ac signal across the three groups of peaks. **d.** Quantification of scATACseq signal at Sox17-bound enhancers in *WT* stem and *Apc^−/−^ Ad-stem* clusters. Dotted lines indicate >2.5 fold gain or loss in accessibility for Sox17-bound loci *Apc ^−/−^Ad-stem* as compared to *WT* stem. Heatmap on the right shows pathways enriched within the genes linked (50 kb distance) to the loci with Sox17 binding and significant accessibility gain as seen in the X-Y plot on the left. **e.** Representative area of adenomas formed in day 14 *Apc ^−/−^*(left) and *Apc ^−/−^ Sox17 ^−/−^* (right) mice from Fig. 4b showing reduction in adenoma size and distribution. **f.** IHC of SOX17in day 14 *Apc ^−/−^ Sox17 ^−/−^* intestinal sections showing adenoma regions that escaped Cre-mediated recombination. Red arrowheads indicate adenomas, and cyan arrowheads show regions with Sox17 expression. **g.** Heatmap showing β-catenin and SOX17 CUT&RUN-seq (CnR) signal at SOX17-independent and -dependent β-catenin targets in presence and absence of Sox17 in adenomas.

**Extended Data Figure 8.**
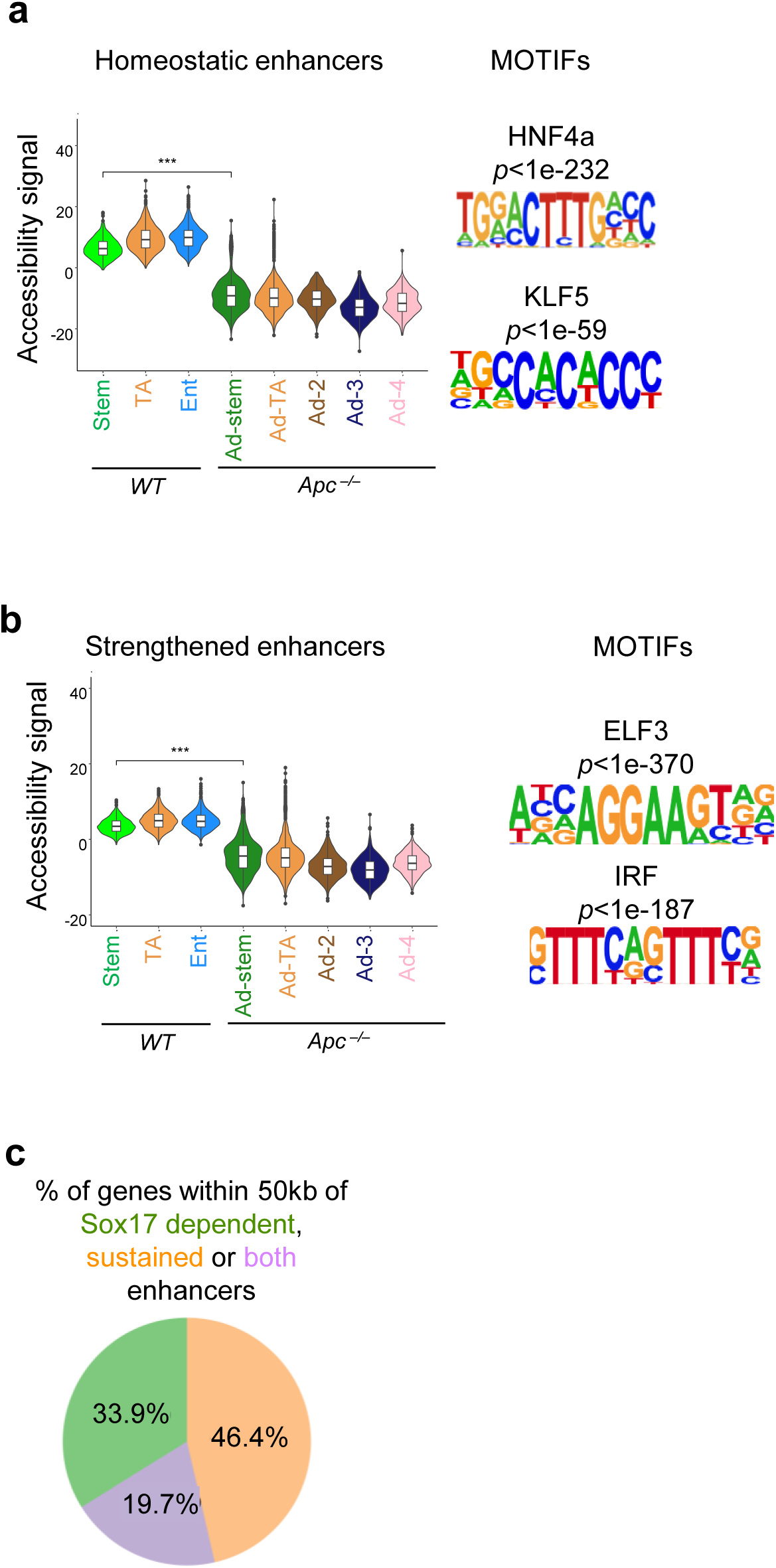
**a.** Violin plots showing accessibility signal at homeostatic enhancers that are predominantly accessible in *WT* clusters. Motifs enriched for these homeostatic enhancers are highlight on the right (JASPAR database). **b.** Violin plot showing accessibility signal at enhancers with strengthened activity in absence of SOX17, as shown in Fig 4a, Motifs enriched for these strengthened enhancers are highlighted on the right. **c.** Pie chart showing percentage of genes linked (within 50 kb) to either *Sox17*-dependent or sustained enhancers, or to both groups.

**Extended Data Figure 9.**
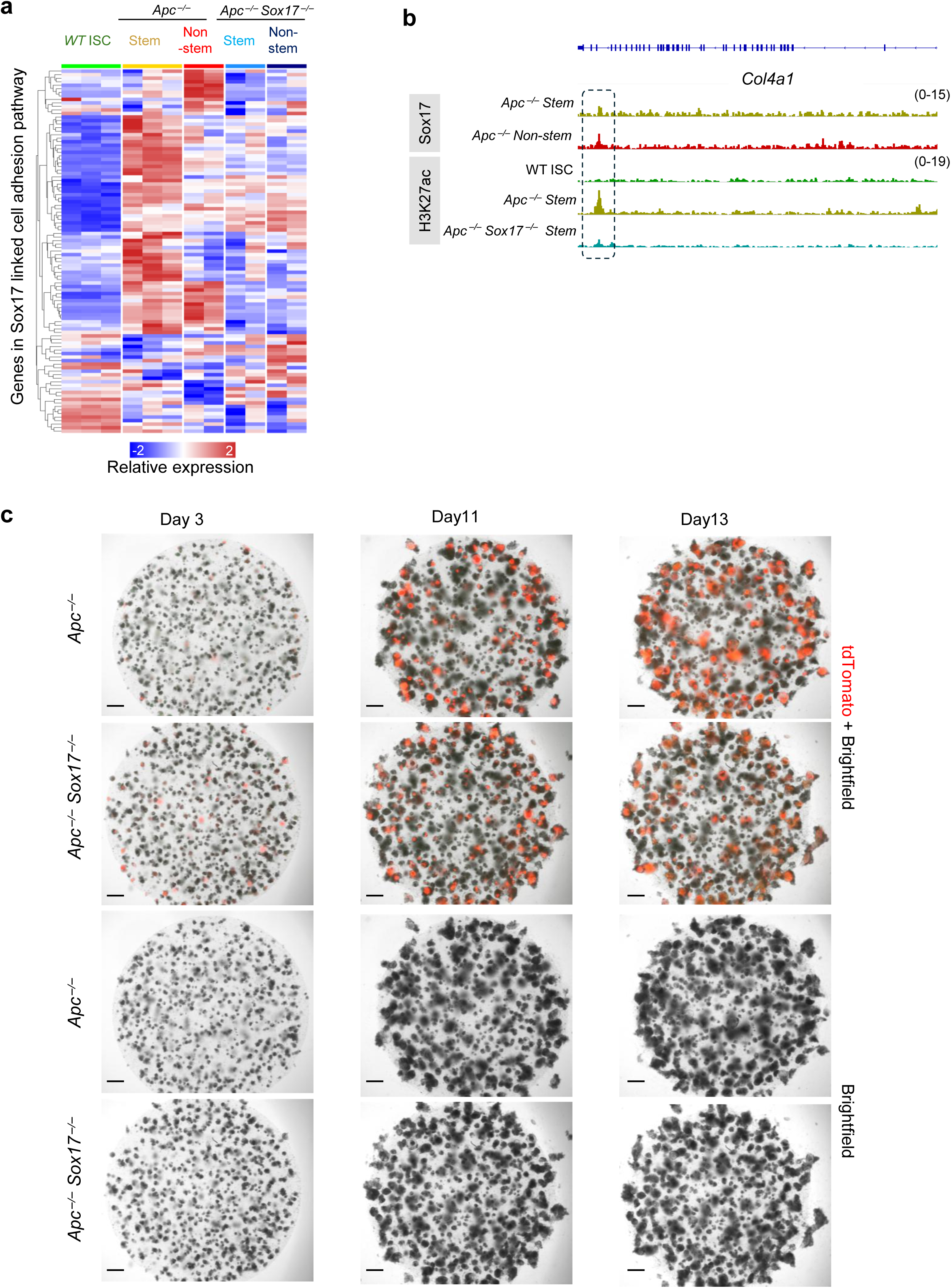
**a.** Heatmap showing relative expression (mRNA-seq, Log2(RPKM+1)) of genes enriched in the cell adhesion and cell motility related pathways from Figure 5a. Hierarchical clustering with “ward.d2” method was applied to identify distinct groups. **b.** Genome viewer tracks around the *Col4a1* locus showing nearby active enhancer (H3K27ac) signals and Sox17 CnR peaks. The dotted box highlights an enhancer bound by Sox17 in *Apc ^−/−^*cells that loses activity in *Apc ^−/−^ Sox17 ^−/−^* cells. **c.** Fluorescent and brightfield images showing progression of *Apc^−/−^*and *Apc^−/−^ Sox17 ^−/−^* organoids imaged across the entire Matrigel dome. Scale bar = 500 μm.

**Extended Data Figure 10.**
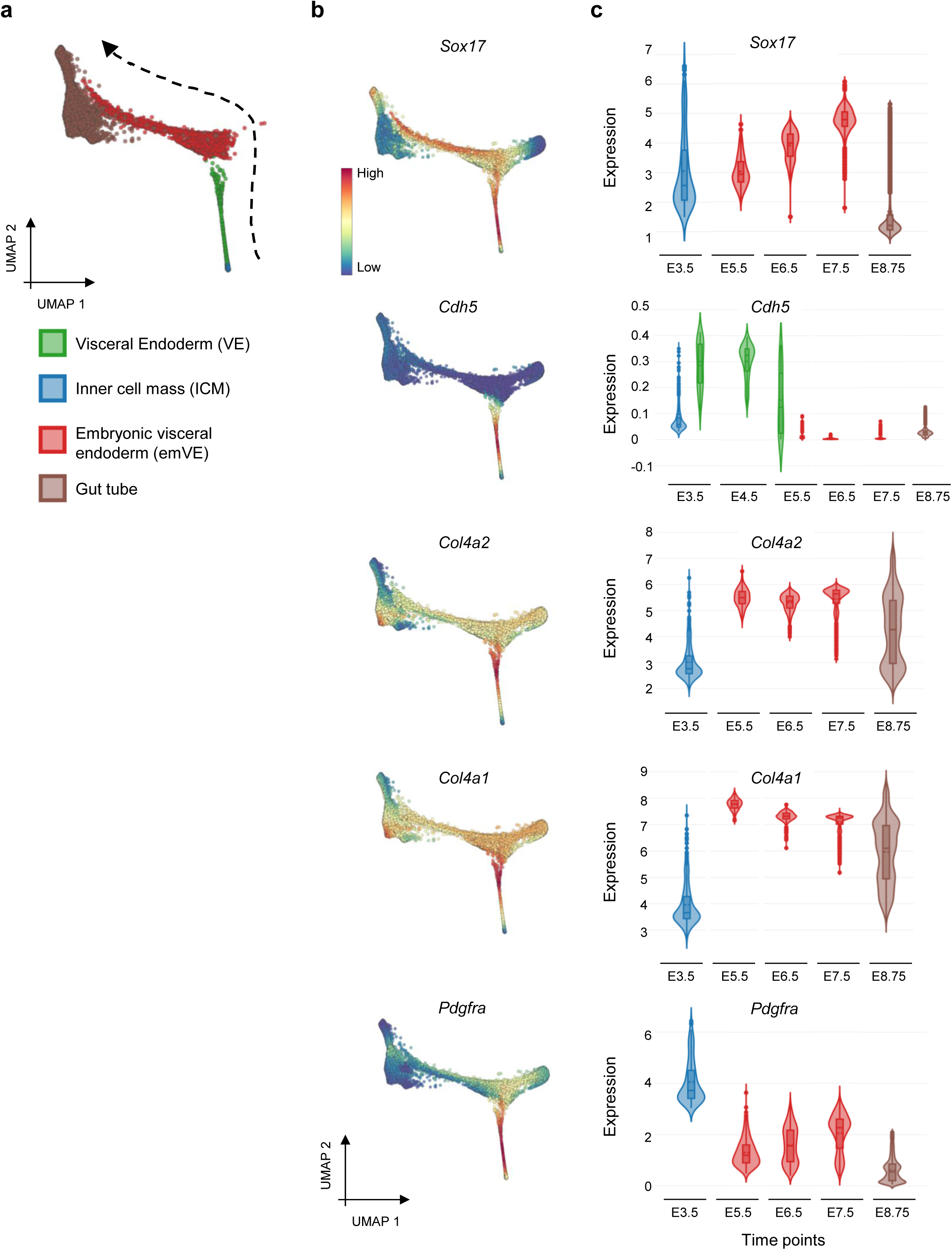
Analysis using data from https://endoderm-explorer.com/ to identify the gene expression patterns in developing mouse endoderm during gut tube formation. **a.** UMAP projection of cells at different stages of endodermal development. The dotted arrow indicates the direction of differentiation from endodermal cells to gut tube cells. **b.** UMAPs showing overlapping expression of *Sox17* and its target genes in adenoma cells (*Cdh5, Col4a2, Col4a1* and *Pdgfra*) across multiple stages of endoderm development and gut tube formation. **c.** Violin plots showing imputed log gene expression of these genes across the cell types present at different timepoints of endodermal development.

## REFERENCES

1. Hu, Z. et al. Quantitative evidence for early metastatic seeding in colorectal cancer. Nat Genet 51, 1113–1122 (2019).

2. Gupta, P.B., Pastushenko, I., Skibinski, A., Blanpain, C. & Kuperwasser, C. Phenotypic Plasticity: Driver of Cancer Initiation, Progression, and Therapy Resistance. Cell Stem Cell 24, 65–78 (2019).

3. Vasquez, E.G. et al. Dynamic and adaptive cancer stem cell population admixture in colorectal neoplasia. Cell Stem Cell 29, 1213–1228.e8 (2022).

4. Househam, J. et al. Phenotypic plasticity and genetic control in colorectal cancer evolution. Nature 611, 744–753 (2022).

5. Dalerba, P. et al. Single-cell dissection of transcriptional heterogeneity in human colon tumors. Nat Biotechnol 29, 1120–7 (2011).

6. He, X. et al. Tumor-initiating stem cell shapes its microenvironment into an immunosuppressive barrier and pro-tumorigenic niche. Cell Rep 36, 109674 (2021).

7. Bala, P. et al. Aberrant cell state plasticity mediated by developmental reprogramming precedes colorectal cancer initiation. Sci Adv 9, eadf0927 (2023).

8. Flavahan, W.A., Gaskell, E. & Bernstein, B.E. Epigenetic plasticity and the hallmarks of cancer. Science 357(2017).

9. Goto, N. et al. SOX17 enables immune evasion of early colorectal adenomas and cancers. Nature 627, 636–645 (2024).

10. Mzoughi, S. et al. Oncofetal reprogramming drives phenotypic plasticity in WNT-dependent colorectal cancer. Nat Genet 57, 402–412 (2025).

11. Becker, W.R. et al. Single-cell analyses define a continuum of cell state and composition changes in the malignant transformation of polyps to colorectal cancer. Nat Genet 54, 985–995 (2022).

12. Powell, S.M. et al. APC mutations occur early during colorectal tumorigenesis. Nature 359, 235–7 (1992).

13. Vermeulen, L. et al. Defining stem cell dynamics in models of intestinal tumor initiation. Science 342, 995–8 (2013).

14. Roerink, S.F. et al. Intra-tumour diversification in colorectal cancer at the single-cell level. Nature 556, 457–462 (2018).

15. Hao, Y. et al. Integrated analysis of multimodal single-cell data. Cell 184, 3573–3587.e29 (2021).

16. Stuart, T., Srivastava, A., Madad, S., Lareau, C.A. & Satija, R. Single-cell chromatin state analysis with Signac. Nat Methods 18, 1333–1341 (2021).

17. Dow, L.E. et al. Apc Restoration Promotes Cellular Differentiation and Reestablishes Crypt Homeostasis in Colorectal Cancer. Cell 161, 1539–1552 (2015).

18. Stancikova, J. et al. NKD1 marks intestinal and liver tumors linked to aberrant Wnt signaling. Cell Signal 27, 245–56 (2015).

19. Ji, W., Zhuang, X., Jiang, W.G. & Martin, T.A. Tight junctional protein family, Claudins in cancer and cancer metastasis. Front Oncol 15, 1596460 (2025).

20. Korinek, V. et al. Constitutive transcriptional activation by a beta-catenin-Tcf complex in APC−/− colon carcinoma. Science 275, 1784–7 (1997).

21. Heino, S. et al. Lef1 restricts ectopic crypt formation and tumor cell growth in intestinal adenomas. Sci Adv 7, eabj0512 (2021).

22. Lu, Z. et al. Characterization of cis-regulatory elements and functional variants in colorectal cancer using epigenomics and CRISPRi screenings. Nat Cancer (2025).

23. Pajtler, K.W. et al. YAP1 subgroup supratentorial ependymoma requires TEAD and nuclear factor I-mediated transcriptional programmes for tumorigenesis. Nat Commun 10, 3914 (2019).

24. Peppino, G. et al. Teneurins: Role in Cancer and Potential Role as Diagnostic Biomarkers and Targets for Therapy. Int J Mol Sci 22(2021).

25. Eide, P.W., Bruun, J., Lothe, R.A. & Sveen, A. CMScaller: an R package for consensus molecular subtyping of colorectal cancer pre-clinical models. Sci Rep 7, 16618 (2017).

26. He, G. et al. Formin-like 2 promotes angiogenesis and metastasis of colorectal cancer by regulating the EGFL6/CKAP4/ERK axis. Cancer Sci 114, 2014–2028 (2023).

27. Huang, D., Du, C., Ji, D., Xi, J. & Gu, J. Overexpression of LAMC2 predicts poor prognosis in colorectal cancer patients and promotes cancer cell proliferation, migration, and invasion. Tumour Biol 39, 1010428317705849 (2017).

28. Du, Y.C. et al. Induction and down-regulation of Sox17 and its possible roles during the course of gastrointestinal tumorigenesis. Gastroenterology 137, 1346–57 (2009).

29. Lee, H.-O. et al. Lineage-dependent gene expression programs influence the immune landscape of colorectal cancer. Nature Genetics 52, 594–603 (2020).

30. Gaynor, L. et al. Crypt density and recruited enhancers underlie intestinal tumour initiation. Nature 640, 231–239 (2025).

31. Chen, B. et al. Differential pre-malignant programs and microenvironment chart distinct paths to malignancy in human colorectal polyps. Cell 184, 6262–6280.e26 (2021).

32. Oyama, T. et al. Further upregulation of beta-catenin/Tcf transcription is involved in the development of macroscopic tumors in the colon of ApcMin/+ mice. Carcinogenesis 29, 666–72 (2008).

33. Kluiver, T.A. et al. Differential Wnt/beta-catenin signaling via TCF7L2/LEF1 binding specificity shapes cellular and tumor phenotypes. Proc Natl Acad Sci U S A 123, e2528450123 (2026).

34. Feng, J., Liu, T., Qin, B., Zhang, Y. & Liu, X.S. Identifying ChIP-seq enrichment using MACS. Nat Protoc 7, 1728–40 (2012).

35. Kazakevych, J., Sayols, S., Messner, B., Krienke, C. & Soshnikova, N. Dynamic changes in chromatin states during specification and differentiation of adult intestinal stem cells. Nucleic Acids Res 45, 5770–5784 (2017).

36. Kanai-Azuma, M. et al. Depletion of definitive gut endoderm in Sox17-null mutant mice. Development 129, 2367–2379 (2002).

37. Engert, S. et al. Wnt/β-catenin signalling regulates Sox17 expression and is essential for organizer and endoderm formation in the mouse. Development 140, 3128–38 (2013).

38. Mukherjee, S. et al. Sox17 and β-catenin co-occupy Wnt-responsive enhancers to govern the endoderm gene regulatory network. Elife 9(2020).

39. Mukherjee, S., Luedeke, D.M., McCoy, L., Iwafuchi, M. & Zorn, A.M. SOX transcription factors direct TCF-independent WNT/β-catenin responsive transcription to govern cell fate in human pluripotent stem cells. Cell Rep 40, 111247 (2022).

40. Sinner, D. et al. Sox17 and Sox4 differentially regulate beta-catenin/T-cell factor activity and proliferation of colon carcinoma cells. Mol Cell Biol 27, 7802–15 (2007).

41. Tian, Y. et al. APC and P53 mutations synergise to create a therapeutic vulnerability to NOTUM inhibition in advanced colorectal cancer. Gut 72, 2294–2306 (2023).

42. Zhang, W. et al. Epigenetic Inactivation of the Canonical Wnt Antagonist SRY-Box Containing Gene 17 in Colorectal Cancer. Cancer Research 68, 2764–2772 (2008).

43. Jadhav, U. et al. Acquired Tissue-Specific Promoter Bivalency Is a Basis for PRC2 Necessity in Adult Cells. Cell 165, 1389–1400 (2016).

44. Goswami, S. & Gao, N. KLF5 Governs Stemness in the Adult Intestinal Stem Cell Niche. Cell Mol Gastroenterol Hepatol 9, 705–706 (2020).

45. Ogden, S. et al. Phenotypic heterogeneity and plasticity in colorectal cancer metastasis. Cell Genom 5, 100881 (2025).

46. Dunbar, K. et al. Aspirin Rescues Wnt-Driven Stem-like Phenotype in Human Intestinal Organoids and Increases the Wnt Antagonist Dickkopf-1. Cell Mol Gastroenterol Hepatol 11, 465–489 (2021).

47. Nowotschin, S. et al. The emergent landscape of the mouse gut endoderm at single-cell resolution. Nature 569, 361–367 (2019).

48. Pennisi, E. How a growth control path takes a wrong turn to cancer. Science 281, 1438–9, 1441 (1998).

49. Laisné, M., Lupien, M. & Vallot, C. Epigenomic heterogeneity as a source of tumour evolution. Nat Rev Cancer 25, 7–26 (2025).

50. de Sousa e Melo, F., et al. A distinct role for Lgr5. Nature 543, 676–680 (2017).

51. Shimokawa, M. et al. Visualization and targeting of LGR5. Nature 545, 187–192 (2017).

52. Fumagalli, A. et al. Plasticity of Lgr5-Negative Cancer Cells Drives Metastasis in Colorectal Cancer. Cell Stem Cell 26, 569–578.e7 (2020).

53. Barker, N. et al. Identification of stem cells in small intestine and colon by marker gene Lgr5. Nature 449, 1003–7 (2007).

54. Colnot, S. et al. Colorectal cancers in a new mouse model of familial adenomatous polyposis: influence of genetic and environmental modifiers. Lab Invest 84, 1619–30 (2004).

55. Sato, T. et al. Single Lgr5 stem cells build crypt-villus structures in vitro without a mesenchymal niche. Nature 459, 262–5 (2009).

56. Lorzadeh, A. et al. Nucleosome Density ChIP-Seq Identifies Distinct Chromatin Modification Signatures Associated with MNase Accessibility. Cell Rep 17, 2112–2124 (2016).

57. Zambanini, G., Nordin, A., Jonasson, M., Pagella, P. & Cantù, C. A new CUT&RUN low volume-urea (LoV-U) protocol optimized for transcriptional co-factors uncovers Wnt/β-catenin tissue-specific genomic targets. Development 149(2022).

58. Buenrostro, J.D., Giresi, P.G., Zaba, L.C., Chang, H.Y. & Greenleaf, W.J. Transposition of native chromatin for fast and sensitive epigenomic profiling of open chromatin, DNA-binding proteins and nucleosome position. Nat Methods 10, 1213–8 (2013).

59. Haber, A.L. et al. A single-cell survey of the small intestinal epithelium. Nature 551, 333–339 (2017).

60. Pinello, L., Farouni, R. & Yuan, G.C. Haystack: systematic analysis of the variation of epigenetic states and cell-type specific regulatory elements. Bioinformatics 34, 1930–1933 (2018).

